# Enhancer AAVs for targeting spinal motor neurons and descending motor pathways in rodents and macaque

**DOI:** 10.1101/2024.07.30.605864

**Authors:** Emily Kussick, Nelson Johansen, Naz Taskin, Ananya Chowdhury, Meagan A. Quinlan, Alex Fraser, Andrew G. Clark, Brooke Wynalda, Refugio Martinez, Erin L. Groce, Melissa Reding, Elizabeth Liang, Lyudmila Shulga, Cindy Huang, Tamara Casper, Michael Clark, Windy Ho, Yuan Gao, Cindy T.J. van Velthoven, Cassandra Sobieski, Rebecca Ferrer, Melissa R. Berg, Britni C. Curtis, Chris English, Jesse C. Day, Michal G. Fortuna, Nicholas Donadio, Dakota Newman, Shenqin Yao, Anish Bhaswanth Chakka, Jeff Goldy, Amy Torkelson, Junitta B. Guzman, Rushil Chakrabarty, Beagen Nguy, Nathan Guilford, Trangthanh H. Pham, Vonn Wright, Kara Ronellenfitch, Robyn Naidoo, Jaimie Kenney, Ali Williford, Charu Ramakrishnan, Antonia Drinnenberg, Kathryn Gudsnuk, Bargavi Thyagarajan, Kimberly A. Smith, Nick Dee, Karl Deisseroth, Hongkui Zeng, Zizhen Yao, Bosiljka Tasic, Boaz P. Levi, Rebecca Hodge, Trygve E. Bakken, Ed S. Lein, Jonathan T. Ting, Tanya L. Daigle

## Abstract

Experimental access to cell types within the mammalian spinal cord is severely limited by the availability of genetic tools. To enable access to lower motor neurons (LMNs) and LMN subtypes, we generated single cell multiome datasets from mouse and macaque spinal cords and discovered putative enhancers for each neuronal population. We cloned these enhancers into adeno-associated viral vectors (AAVs) driving a reporter fluorophore and functionally screened them in mouse. We extensively characterized the most promising candidate enhancers in rat and macaque and developed an optimized pan LMN enhancer virus. Additionally, we generated derivative viruses expressing iCre297T recombinase or ChR2-EYFP for labeling and functional studies, and we created a single vector with combined enhancer elements to achieve simultaneous labeling of layer 5 extratelencephalic projecting (ET) neurons and LMNs. This unprecedented LMN toolkit will enable future investigations of cell type function across species and potential therapeutic interventions for human neurodegenerative diseases.

## Introduction

The spinal cord is a vital part of the central nervous system (CNS) and functions to relay information back and forth between the brain and the body. It is comprised of a diversity of cell types that have been classically defined by their morphological and physiological properties, and more recently by their single cell transcriptomes^1–6^. Lower motor neurons (LMNs) are specialized types of cholinergic cells that reside in the brainstem and the ventral horn of the spinal cord. They receive input from L5 ET neurons located in the brain directly in humans or indirectly via an intermediate synapse in rodents and in turn, initiate actions by controlling effector muscles in the periphery^7^. Degeneration of motor neurons (MNs) can profoundly impact the initiation and control of movement and is a hallmark of many devastating diseases such as Amyotrophic Lateral Sclerosis (ALS) and Spinal Muscular Atrophy (SMA) for which there are no known cures^8,9^.

While considerable progress has been made on LMN cell type definitions, there is still much to learn about their overall organization and interconnectivity within the spinal cord and with cells throughout the body in both healthy and pathological conditions. Experimental access to LMNs in rodents still largely depends on the Chat-IRES-Cre line which broadly labels all cholinergic neurons throughout the body^10^ or vectors containing the homeobox gene *Hb9* derived promoter fragments^11–14^ which exhibit varying levels of specificity and strength depending on the experimental context. Importantly, no genetic tools exist to selectively target LMNs across species or to target any LMN subtypes (i.e. alpha, beta, or gamma) in any species or for simultaneous and selective targeting of LMNs and L5 ET neurons. This precludes the possibility of refined targeting strategies, such as those needed to better define cell type function and for potential therapeutic interventions.

We previously developed an approach to identify functional enhancer elements within the genome and used it to create adeno associated viruses (AAVs) to target specific interneuron and excitatory projection neuron subtypes within the mouse neocortex and to a limited degree in the non-human primate brain^15,16^. More recently, we scaled and expanded the effort to create larger suites of viral genetic tools for near complete coverage of the neocortical neuron landscape and of major non-neuronal types in mouse^17,18^. These successful efforts showed generalizability of the enhancer AAV technology platform and suggested the potential for future tool development outside of the brain.

Here, we provide high-quality, single-cell multiome datasets from the mouse and macaque spinal cord to reveal genome-wide regions of open chromatin across major cell classes. We use these datasets to discover putative enhancer elements for LMNs and LMN subtypes with potential for cross-species conservation and functionally test all of them in mouse and a subset of the most promising in rat and macaque using clinically relevant routes of administration to show they effectively target the homologous cell populations. We identify the core functional region of the top pan LMN enhancer and create an optimized virus. We also generate and validate enhancer AAVs driving either iCre297T or ChR2-EYFP2 in LMN populations in rodents. Additionally, we demonstrate that enhancers targeting both L5 ET and LMN populations can be stitched in *cis* in a single viral vector to effectively achieve the combined labeling pattern within the CNS of rodents.

## Results

### Generation of 10X Multiome data from mouse and macaque spinal cord and identification of major cell classes

To identify LMN-specific enhancers, we isolated nuclei from cervical and lumbar segments of spinal cord in mouse and macaque. We selectively enriched for neurons using fluorescence-activated nuclei sorting (FANS) to isolate nuclei at a ratio of 70% NeuN+ and 20% OLIG2-, 10% OLIG2+ (see **Methods Details**). Gene expression and open chromatin were profiled from individual nuclei using 10X Multiome. In total, we collected high quality data from 73,120 single nuclei from 5 mice and 34,344 single nuclei from 4 macaques (**Figure 1A**). High quality nuclei were retained that expressed more than 1000 genes, had less than 3% mitochondrial reads and doublet scores less than 0.3, and had ATAC-seq fragments in transcription start sites and that spanned nucleosomes based on standards reported by ENCODE^19^.

**Figure 1.**
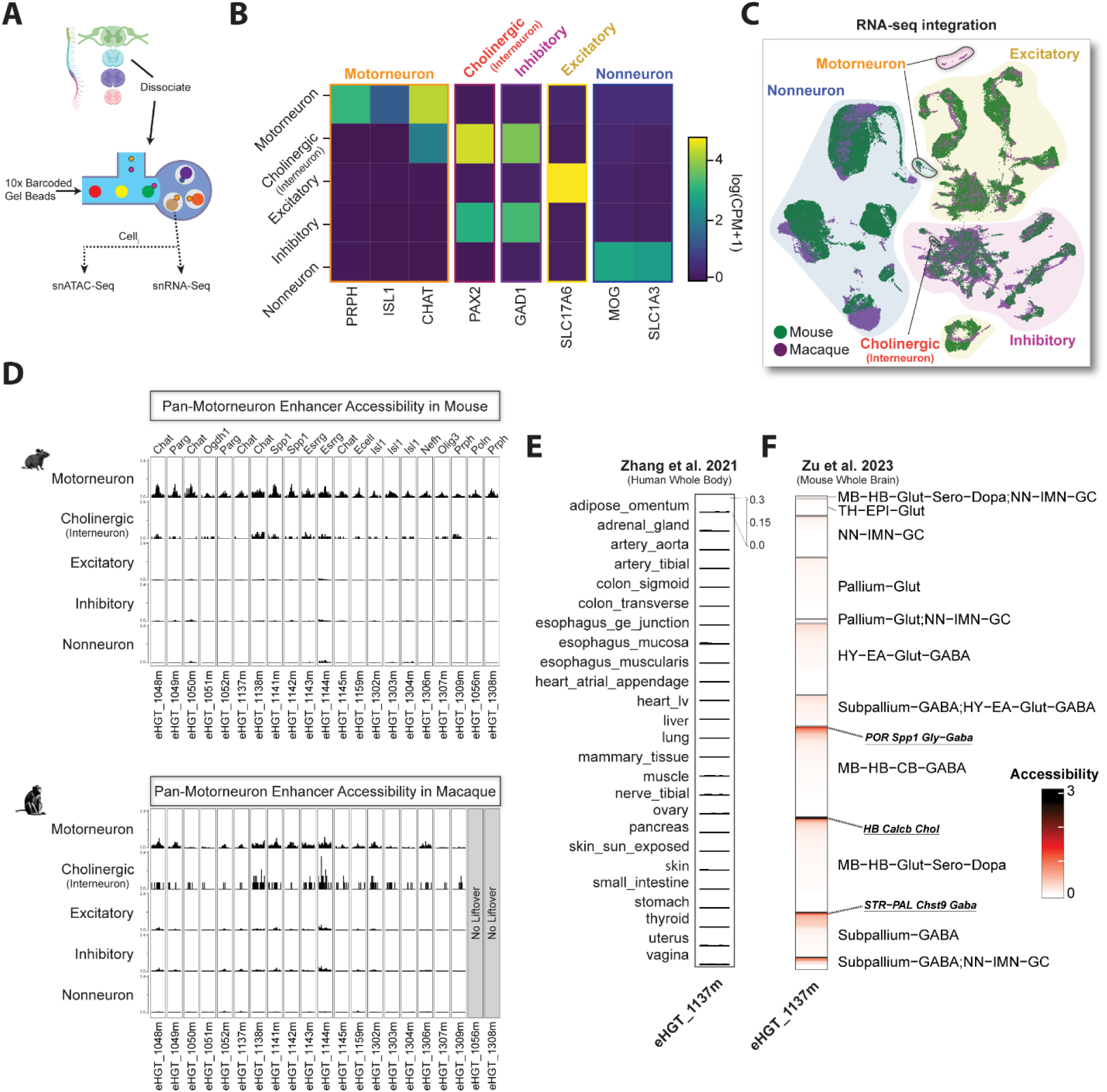
Generation of 10X multiome datasets from mouse and macaque spinal cord and motor neuron enhancer discovery. (**A**) Schematic of 10X Multiome profiling of spinal cord. (**B**) Heatmap of log2(CPM+1) expression of spinal cord marker genes. Colored boxes group the marker genes by broad spinal cord cell types. (**C**) Uniform manifold approximation and projection (UMAP) dimensional reduction of integrated snRNA-seq spinal cord data from Mouse and Macaque. (**D**) Chromatin accessibility of pan-motor neuron enhancers in both Mouse and Macaque. (**E**) Chromatin accessibility of pan-motor neuron enhancer eHGT_1137m in multiple non-brain tissues across the human body (Zhang et al. 2021) (**F**) Chromatin accessibility of pan-motor neuron enhancer eHGT_1137m across the mouse whole brain (Li et al. 2023). Each row indicates the normalized accessibility of eHGT_1137m within each subclass (n=318) grouped by neighborhoods (n=11) from Yao et al. 2023. Subclasses in which significant expression was observed are highlighted in bold and underlined text. MB=midbrain, HB=hindbrain, TH=thalamus, EPI=epithalamus, HY=hypothalamus, CB=cerebellum.

Spinal cord cell types were defined based on iterative clustering of transcriptomic data, and all types had distinct marker genes (see **Methods Details**;^1,2^). Cell type identities were assigned based on enriched expression of known marker genes for LMNs (CHAT, ISL1, PRPH), other excitatory neurons (SLC17A7), inhibitory neurons (GAD1), cholinergic interneurons (CHAT, GAD1, PAX2), and non-neuronal types (SLC1A3, MOG) (**Figure 1B**). Macaque and mouse snRNA-seq data was aligned using scVI^20^. We found consistent coverage of a conserved set of cell types across species (**Figure 1C**) highlighting conservation of distinct marker genes for spinal cord cell types.

### Identification of putative pan motor neuron enhancers and deeper analysis of the 1137m enhancer

To identify putative enhancers specific to LMNs, we aggregated the snATAC-seq data for all nuclei within each species and cell class and identified peaks specific to mouse LMNs that were proximal to published LMN marker genes^1,2^ (**Figure 1D**). Many pan-LMN enhancers had conserved accessibility between mouse and macaque (**Figure 1D**). We further evaluated cell type- and tissue-specificity of each pan-LMN enhancer in human non-brain tissues^21^ and in the whole mouse brain^22^ by assessing the accessibility of the corresponding genomic region (**Table S1)**. 1137m was found to be highly specific to LMNs and inaccessible in all non-brain tissues (see **Methods Details**; **Figure 1E**). Additionally, based on mouse whole brain ATAC profiling, we found accessibility was limited to three rare cell types (HB Calcb Chol, POR Spp1 Gly-Gaba and STR-PAL Chst9 Gaba) (see region definitions in **Figure legend 1F**) that are predominantly found in brainstem (**Figure 1F**). Interestingly, only HB Calcb Chol expresses the enhancer proximal LMN marker *Chat*, suggesting that 1137m has altered regulatory function in the other two cell types. We also analyzed accessibility proximal to previously described^5,11,23^ motor neuron marker genes from early development (Mnx1, Phox2b, Lhx3, Slit2 and Slit3) and found highly specific accessibility only proximal to Mnx1 (**Figure S1**). These data suggest possible divergence in the regulatory landscape in LMNs from early development into adulthood.

### Generation of enhancer-driven fluorophore AAVs for pan spinal motor neuron labeling across species

We first assessed the ability of reported blood-brain-barrier (BBB) permeant capsids, PHP.eB and 9P31^24,25^ to effectively cross the blood-spinal cord barrier (BSCB) and transduce neurons broadly within the spinal cord following retro-orbital (RO) delivery to mice. We packaged recombinant AAVs (rAAVs) containing a pan neuronal human synapsin 1 (hSyn1) promoter driving a histone 2B (H2B) tethered directly to a SYFP2 fluorescent protein for nuclear enrichment called CN1839^15^ and delivered 5.0 × 10^11^ GC of PHP.eB or 9P31-serotyped CN1839 viruses to wild-type mice and assessed viral expression in brain and spinal cord 3-4 weeks post-infection (**Figure S2A, B**). We found that both serotypes exhibited strong nuclear SYFP2 neuronal labeling throughout the brain and in all levels of spinal cord (**Figure S2A, B**). For comparison, we also tested PHP.S serotyped virus, a broad peripheral nervous system (PNS) directed capsid with some reported spinal cord expression^24^, and found the same viral dose gave very little expression in brain or spinal cord following RO administration (**Figure S2A, B**). Subsequent anti-Chat immunostaining of 9P31 or PHP.eB CN1839 cervical spinal cord sections revealed little co-labeling of LMNs within ventral horn regions (33.5%-40.6%) and predominate (100%) labeling of partition cells (PCs) following RO injection (**Figure S2C-E**). Weak LMN labeling likely reflected hSyn1 promoter activity, not capsid tropism, as this promoter has been found to poorly label cholinergic neurons throughout the CNS (data not shown). Collectively, these data suggested that either PHP.eB or 9P31 capsid would work for non-invasive screening of putative enhancers targeting spinal cord cell types, and therefore we selected the PHP.eB capsid because we have used it extensively in our previous work^15,16^.

To functionally test the putative enhancers for LMNs, we cloned them into an rAAV genome upstream of a minimal β-globin promoter driving the SYFP2 fluorescent protein, delivered 5.0 × 10^11^ GC of PHP.eB viruses RO to wild-type mice, and analyzed expression in spinal cord 3-4 weeks post-infection (see experimental workflow in **Figure 2A**). We functionally screened a total of 21 putative LMN enhancers *in vivo* and found that 16 of the enhancer-containing AAVs (76%) drove SYFP2+ cell body labeling within the transverse spinal cord section, 13 of these showed SYFP2+ labeling of cells within the ventral horn (61%) and 5 showed no SYFP2+ expression (24% failure rate) in the sections analyzed (**Figure S2 A, B**). The HCT1-1137m-SYFP2 enhancer virus (referred to as HCT1 from here forward) appeared initially to exhibit the strongest labeling of spinal motor neurons (**Figure 2B, and Figure S2A**) and therefore, we selected it for additional examination using a variety of experimental techniques. We performed light sheet imaging of cleared spinal cords from HCT1 virus infected animals and compared the expression directly to that achieved with a ubiquitous promoter-driven SYFP2 PHP.eB virus (CN4358 CAG-SYFP2) (**Figure 2C** and **Videos S1, 2**). Here we found markedly more selective SYFP2 expression within the ventral horn at all levels of spinal cord for HCT1 compared to CN4358 (**Figure 2C and Video S1, 2**). To confirm that the labeled cells were positive for choline acetyltransferase (Chat), we performed anti-Chat immunostaining on fixed spinal cord sections from RO-injected HCT1 animals and evaluated co-localization of native SYFP2 fluorescence with Chat immunoreactivity throughout the spinal cord (**Figure 2D**). We identified Chat+ neurons within medial and lateral motor columns (MMC and LMN, respectively), the preganglionic motor column (PGC), phrenic motor neurons of lamina 9 (Ph9), or PCs in lamina 7 in the appropriate anatomical levels based on the Allen Institute spinal cord reference atlas (**Figure 2E, F**). Quantitative analysis of co-localization revealed that the HCT1 virus labelled Chat+ neurons with high specificity in MMC (79.8-90.0%), LMC (90.9-93.4%), and Ph9 (81.2%) and with high completeness of labeling in the same areas (84.6-95.1%) (**Figure 2 G, H and Figure S7A, B**). A complimentary analysis of HCT1 viral labeling of Chat+ cells was also performed using Chat-IRES-Cre;Ai75 tdTomato reporter animals and a coarser anatomical segmentation, which revealed similarly high specificity and completeness of labeling of LMNs in ventral horn (**Figure S6**). In contrast, we observed very little (14.2-25%) viral labelling in partition cells and in neurons within PGC (**Figure 2G, H**). Lastly, we evaluated HCT1-driven viral labelling of Chat+ neurons following two different routes of administration: intracerebroventricular (ICV) and intraspinal (IS) (**Figure 2D, G, H** and **Figure S4C**). Similarly to the RO route, ICV injection of HCT1 resulted in very high specificity and completeness of labeling in MMC, LMC, and Ph9 (90.8-100%; 80.1-85.5%) and sparse labeling of PCs (0%; 0%) and neurons within PGC (0-16.6%; 0-2.0%) (**Figure 2G, H**). Intraspinal injection of HCT1 resulted in enriched LMN labeling relative to CN1839 but considerable off-target labeling in the lateral and ventral funiculus (lf and vf) and the lateral spinal nucleus (LSp) at both conditions tested (**Figure S4C**).

**Figure 2.**
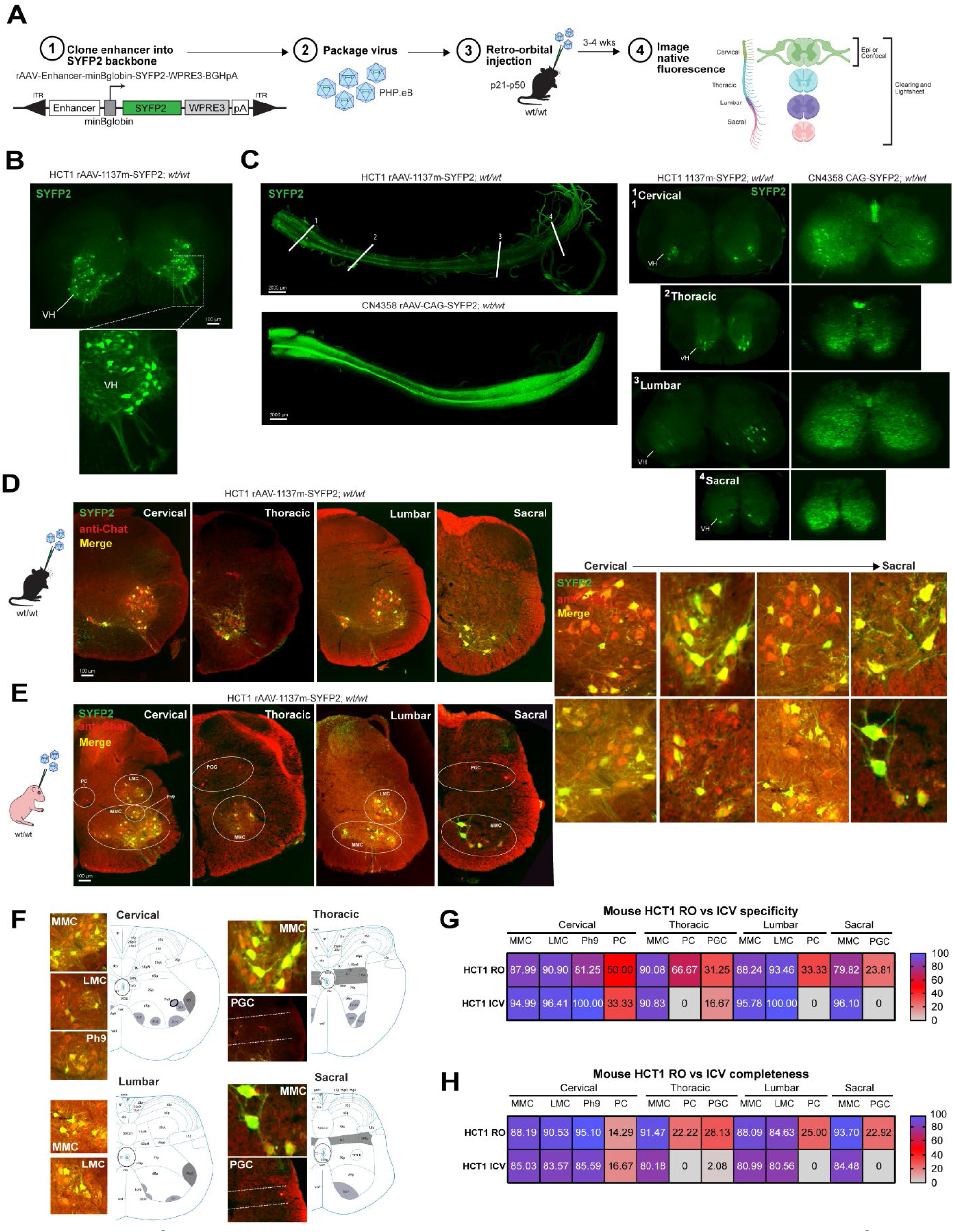
The HCT1 enhancer AAV selectively labels motor neurons in all levels of mouse spinal cord. (**A**) Experimental workflow for enhancer AAV generation and primary screening in mouse. Putative enhancers are cloned into a standard rAAV backbone containing the indicated components. minBglobin = minimal beta-globin promoter, SYFP2 = super yellow fluorescent protein 2, WPRE3 = shortened woodchuck hepatitis virus posttranscriptional regulatory element, pA = bovine growth hormone poly A, ITR = inverted terminal repeats. 5.0 × 10^11^ GC of PHP.eB serotyped virus was delivered via retroorbital (RO) route of administration and spinal cord harvested 3–4-week post-injection and analyzed (**B**) Representative epifluorescence HCT1 screening images of 1137m-driven native SYFP2 fluorescence in 50µm thick transverse spinal cord sections. VH = ventral horn and dashed box denotes area of interest highlighted in the bottom panel. (**C**) Light sheet images of intact mouse spinal cords (left) and 1.2µm thick transverse spinal cord sections from each level of cord (right) that show native SYFP2 expression driven by RO injection of 5.0 × 10^11^ GC of PHP.eB serotyped HCT1 or CN4358 viruses.(**D-E**) Representative high and low magnification epifluorescence images of native SYFP2 fluorescence (green) and anti-Chat immunostaining (red) in 30µm sections from all levels of cord prepared from HCT1 RO-injected (**D**) or ICV-injected mice. (**E**) Anatomical areas of interest or previously described cell types are denoted and labelled in white. MMC = medial motor column, LMC = lateral motor column, Ph9 = phrenic motor neurons of lamina 9, PGC = preganglionic motor column, and PC = partition cells. (**F**) Representative images of HCT1 viral labeling in areas and cell types indicated (left panels). Allen Institute adult mouse spinal cord atlas images are presented in right panels and shaded (grey) to denote corresponding anatomical areas of interest. (**G-H**) Heatmap representation of percent (%) specificity and completeness of HCT1 viral labeling of Chat+ motor neurons in the spinal cord following RO or ICV injection. Legend with scale denotes values ranging from 0 (grey) – 100% (purple). Anatomical areas listed below heatmap do not exist at that level of cord. Percentage values are the average mean value from n = 3 animals and 3 sections per animal per level of cord analyzed.

To determine if the HCT1 virus could label LMNs in rats, we delivered ∼1.50-2.00 × 10^11^ GC of it to rat neonatal pups via ICV injection and analyzed expression by light sheet imaging 3-4 weeks post-infection (**Figure 3A**). Like in mouse, we found that the HCT1 virus selectively labeled neurons within the ventral horn at all levels of spinal cord compared to the CN1839 virus (**Figure 3A and Videos S3,4**). We performed anti-Chat immunolabeling of fixed HCT1 and CN1839 sections and quantified SYFP2 fluorescence and Chat signal co-localization at multiple levels of spinal cord in the same anatomical regions and neuronal types as in mouse (**Figure 3B-F**). This analysis revealed extremely high specificity of the HCT1 virus for Chat+ neurons in MMC, LMC, and Ph9 (95.7-100%), completeness of labeling ranging from 54.6% (LMC, cervical) to 100% (Ph9, cervical), and a complete absence of labeling of PCs (**Figure 3E, F**). CN1839 viral labeling of Chat+ neurons resulted in similar completeness of labeling in MMC, LMC, and in PCs (38.8-80.5%), but less labeling in Ph9 (0%) and overall reduced specificity as expected for a pan-neuronal virus (**Figure 3E, F**). Comparative analysis between mouse and rat identified significant conservation (phylop p-value 0.0388) of the DNA sequence underlying HCT1 which further supports the conserved expression pattern across species.

**Figure 3.**
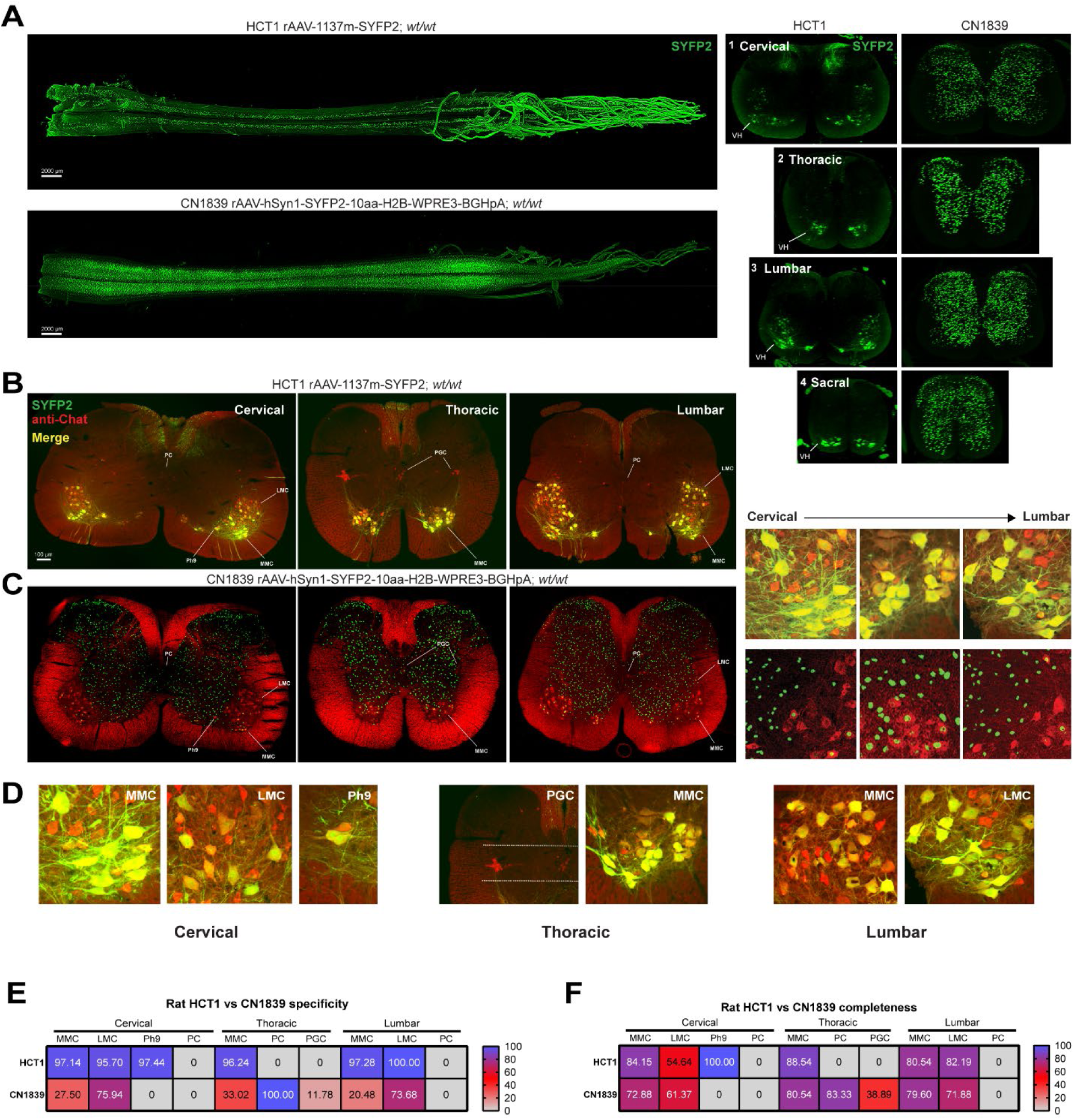
Conserved labeling of spinal motor neurons by the HCT1 enhancer AAV in rat. (**A**) Light sheet images of native SYFP2 fluorescence in intact rat spinal cords (left) and 1.2µm thick transverse spinal cord sections from each level of cord (right) from HCT1or CN1839 infected rats. VH = ventral horn. (**B-C**) Representative confocal images of native SYFP2 (green) and anti-Chat immunostaining (red) in 30µm thick transverse spinal cord sections at the indicated level from ICV injections of HCT1 (B) or CN1839 (C) in rats. Higher magnification images of the ventral horn in each area are also shown (right). Anatomical areas of interest or previously described cell types are denoted and labelled in white. MMC = medial motor column, LMC = lateral motor column, PGC = preganglionic motor column, and PC = partition cells. N = 2 animals per group and 3-6 sections analyzed per animal. (**D**) Representative higher magnification images of HCT1 viral labeling in anatomical areas of interest or previously described cell types. (**E-F**) Heatmap representation of percent (%) specificity and completeness of HCT1 or CN1839 viral labeling of Chat+ motor neurons in anatomical areas and previously defined cell types following ICV injection. Legend with scale denotes values ranging from 0 (grey) – 100% (purple). Percentage values are the average mean value from n = 1 animal and a minimum of 3 sections per animal per level of cord analyzed.

We next tested whether the HCT1-1137m enhancer would exhibit similar activity in LMNs in the non-human primate (NHP) spinal cord following two different clinically relevant routes of administration. We first constructed a vector containing the 1137m enhancer driving a 3XFlag epitope tagged mTFP1 fluorophore (referred to as HCT69 from here forward) and packaged into purified PHP.eB virus (**Figure 4A**). Naïve juvenile macaques were injected with 1.0 × 10^13^ vg/kg of HCT69 virus via the intra-cisterna magna (ICM) or intrathecal (IT) routes of administration and sacrificed 5-7 weeks post-injection for histological analyses (**Figure 4A**; also see **Methods Details**). We then fixed transverse sections throughout the spinal cord from each animal, evaluated native mTFP1 fluorescence (data not shown), and stained with anti-Flag epitope and anti-Chat antibodies to determine the extent of viral labeling in Chat+ neurons in the indicated anatomical regions and in neuronal types (**Figure 4B-D**). ICM administration resulted in highly specific viral labeling of Chat+ neurons in MMC cervical through lumbar cord (85.8-96.3% range), LMC (87.7-93.0%), Ph9 (93.9%) and PGC at sacral level (95.2%) (**Figure 4E**). IT administration resulted in highly specific viral labeling in MMC lumbar and thoracic levels (98.0 and 97.4%, respectively), LMC at lumbar level (95.9%), Ph9 (95.5%) and PGC at thoracic level (87.6%) (**Figure 4E**). No viral labeling was observed in PCs at any level of spinal cord by either route of administration, and completeness of labeling ranged from 0% (PC all levels) to 96.6% (Ph9 cervical level) (**Figure 4E, F**). Collectively, these data demonstrate strong conservation of 1137m enhancer activity in spinal motor neurons throughout the cord and suggest both routes of administration can effectively transduce this neuronal population in macaque.

**Figure 4.**
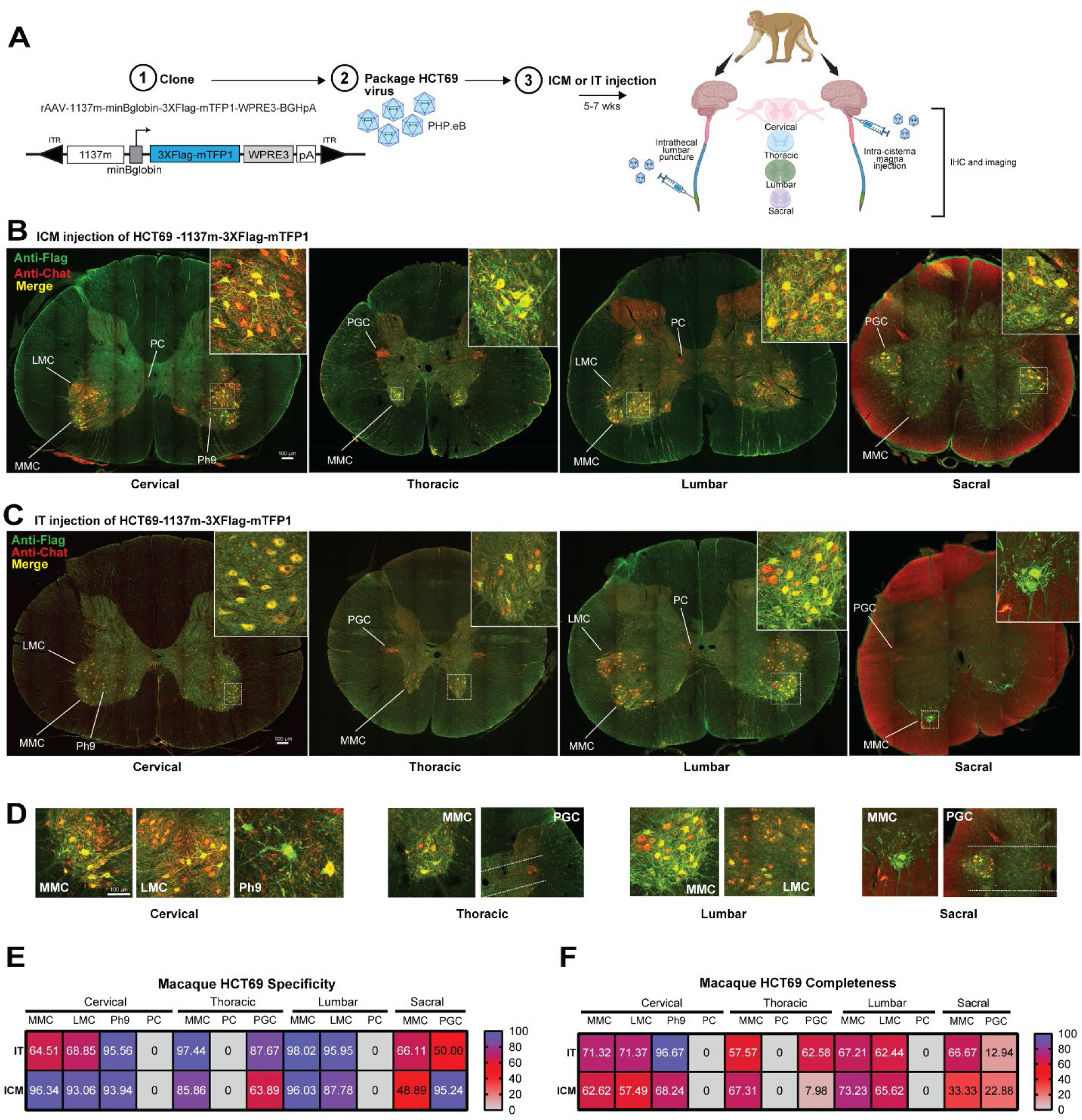
Conserved 1137m enhancer driven labeling of spinal motor neurons in macaque following clinically relevant routes of administration. (**A**) Experimental workflow for HCT69 enhancer AAV generation and testing in macaque by intra-cisternal magna (ICM) or intrathecal (IT) routes of administration. A 3XFlag-mTFP1 gene was inserted in place of SYFP2 in the standard rAAV backbone and packaged in PHP.eB serotype. Purified virus was delivered ICM or IT to young adult macaque animals and spinal cord and other organs were harvested 5-7 weeks post-injection. (**B-C**) Representative images of anti-Flag signal (green) and anti-Chat signal (red) in 30µm thick transverse spinal cord sections at the level indicated prepared from ICM (**B**) or IT (**C**) HCT69-injected animal. Insets show higher magnification view of cells highlighted in the dashed boxed region in each low magnification image. Anatomical areas of interest or previously described cell types are denoted and labelled in white. MMC = medial motor column, LMC = lateral motor column, PGC = preganglionic motor column, PC = partition cells, and Ph9 = phrenic motor neurons of lamina 9. (**E-F**) Heatmap representation of percent (%) specificity and completeness of HCT69 viral labeling of Chat+ motor neurons in anatomical areas and previously defined cell types following IT or ICM injection. Legend with scale denotes values ranging from 0 (grey) – 100% (purple). Percentage values are the average mean value from n = 1 animal and a minimum of 3 sections per animal per level of cord analyzed.

We further evaluated HCT1 and HCT69 viral expression in fixed sections prepared from the brain and peripheral tissue (heart, liver, retina, and dorsal root ganglion (DRG) for mouse; heart, liver, kidney, pancreas, muscle, and DRG for NHP) of RO, ICM, and IT injected animals and directly compared labeling to that achieved in CN1839 injected animals (RO for mouse and ICM for NHP) given the same dose (**Figure S4A and Figure S5)**. In mouse, RO injection of HCT1 resulted in viral labeling in the brain that was mostly confined to hindbrain structures (i.e. deep cerebellar nucleus and various cholinergic brainstem nuclei) and very sparse (liver) to no (heart, retina, DRG) expression in the other organs analyzed (**Figure S4A and Table S3**). ICV injection of HCT1 resulted in no notable expression in heart, liver or brain (**Figure S4B**). By comparison, the RO delivery of the CN1839 virus in mouse resulted in no expression in heart or liver and strong, widespread expression throughout brain and retina (**Figure S4A**). In NHP, the HCT69 virus showed no expression in the brain, or any peripheral organ evaluated with either delivery route, including DRGs that were kept intact and visualized from the ICM cord (**Figure S5B, C and Table S3**). By comparison, NHP sections from CN1839 virus ICM injected animals gave moderate neuronal labeling throughout the brain, with notable pockets of expression in cortical regions (**Figure S5A and Table S3**), consistent with previous reports of AAV9 based virus infectivity ^26,27^. Additionally, we found more widespread neuronal labeling in the cervical spinal cord of the CN1839 animal and observed little (heart) to no expression (all other organs) in the periphery (**Figure S5A and Table S3**). IHC runs were performed on CNS and peripheral organ NHP sections in parallel to rule out any methodological issues that could account for the differences observed. A summary of the biodistribution data for each virus tested by different routes of administration in the study can be found in **Table S3**. These data collectively demonstrate that HCT1 and HCT69 enhancer viruses label LMNs within the body incredibly more selectively than a hSyn1 promoter driven virus and are consistent with the predicted body-wide accessibility profile for the 1137m enhancer in human and mouse epigenomic data (**Figure 1E, F**).

### Generation of optimized pan LMN virus

Higher transgene levels in LMNs may be desirable for functional experiments that require strong effector gene expression and/or to support diverse therapeutic strategies. Hence, we developed an optimized pan LMN virus by first bashing the full-length 1137m enhancer sequence to identify the functional core domain (**Figure 5A**). We cloned 240-243bp 1137m enhancer fragments upstream of the minimal promoter and SYFP2 in the rAAV backbone, packaged PHP.eB serotyped viruses for each (HCT52, HCT53, and HCT54; **Figure 5A**) and screened viruses by RO injection of wild-type mice (**Figure 5B**). The middle 240bp fragment termed Frag2 was found to drive expression in LMNs, which was confirmed with anti-Chat immunostaining (**Figure 5B, C**). To enhance expression in LMNs, we generated a three times Frag2 (3XFrag2) concatemer, cloned it into the SYFP2 expressing vector (termed HCT47), and evaluated native SYFP2 signal alone or with anti-Chat signal by epifluorescence and light sheet imaging in RO injected animals (**Figure 5D-F**). We found that the HCT47 virus strongly labelled LMNs throughout the spinal cord and performed similarly to the HCT1 virus in terms of specificity and completeness of labeling, with the coarser segmentation method, in the ventral horn at the cervical level (HCT47 = 77.8% specific and 83.1% complete; HCT1 = 83.8% specific and 72.35% complete; **Figure 5F-H**). Additionally, we observed an absence of viral labeling in attached DRGs (**Figure 5F and Figure S4, and Table S3**). To determine if vector optimization increased SYFP2 protein levels in spinal cord, we performed western blot analyses on tissue lysates prepared from RO injected HCT47 or HCT1 animals (**Figure 5I**). Here we found a 4.3-fold increase in HCT47-driven levels compared to HCT1-driven reporter protein levels (**Figure 5I, J**). These data demonstrate that concatenation of the 1137m core region significantly increased (p = 0.049) reporter gene expression, while largely maintaining LMN specificity.

**Figure 5.**
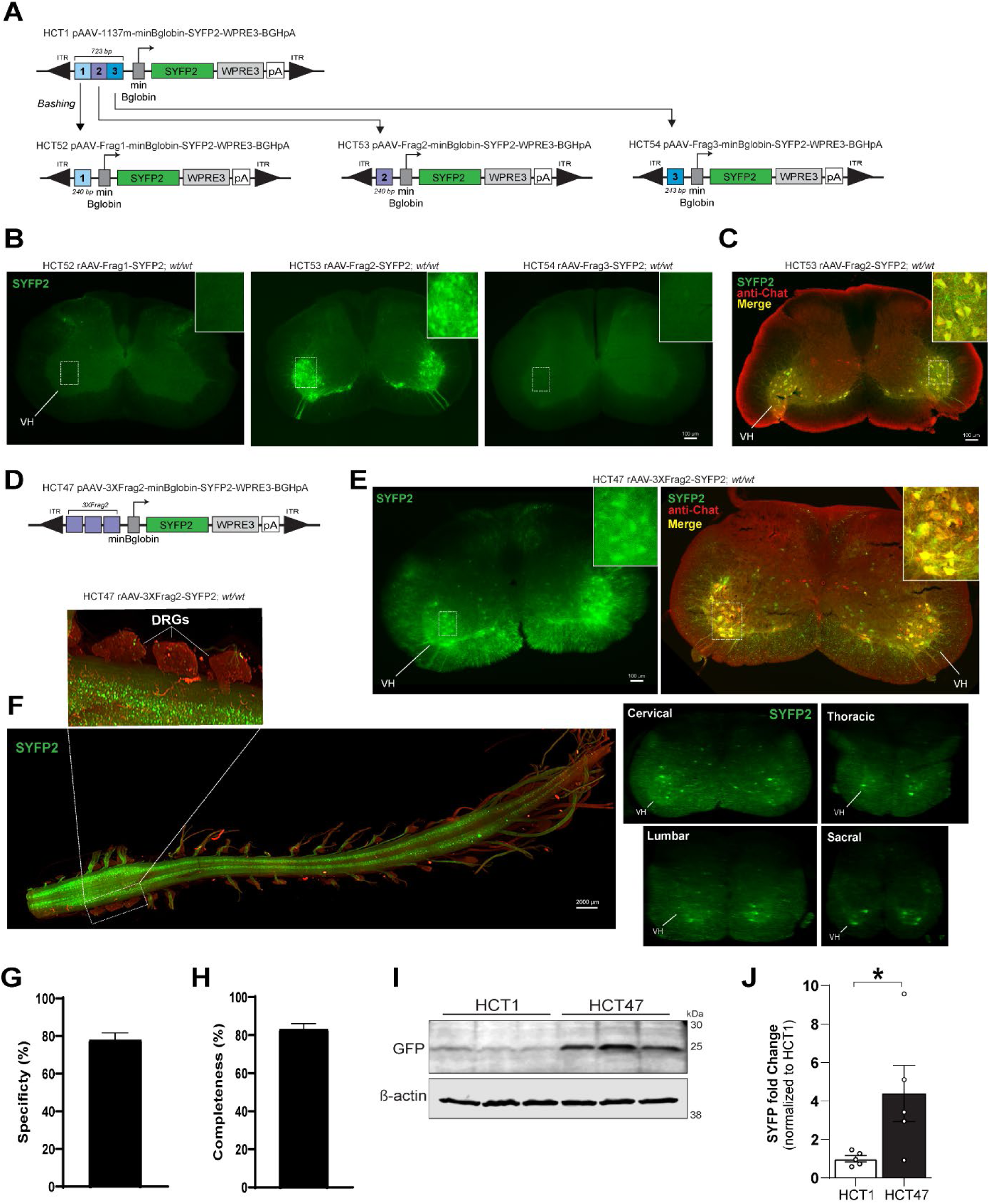
Generation and validation of optimized pan spinal motor neuron enhancer virus. (**A**) Schematic diagram of full-length 1137m enhancer bashing approach and creation of HCT52, HCT53, and HCT54 vectors. (**B**) Representative epifluorescence images of native SYFP2 fluorescence in 50µm thick transverse sections from cervical spinal cord of RO injected HCT52-, HCT53-, or HCT54-animals. (**C**) Representative image of native SYFP2 fluorescence (green) and anti-Chat signal (red) in a 30µm thick transverse cervical spinal cord section from an HCT53 animal. (**D**) Optimized HCT47 vector schematic. Each purple box = fragment 2 (Frag2) from HCT53 cloned in tandem to create the 3XFrag2 concatemer. (**E**) Representative epifluorescence image of native SYFP2 fluorescence in a 50µm thick transverse cervical cord section from a mouse RO injected with the HCT47 virus (left). Right panel shows a representative image of native SYFP2 fluorescence (green) and anti-Chat signal (red) in an HCT47 cervical spinal cord section. Insets in all panels show higher magnification view of cells highlighted in the dashed boxed region in each low magnification image and VH = ventral horn. (**F**) Light sheet images of native SYFP2 fluorescence in intact mouse spinal cord (left) or in 1.2µm thick transverse sections from each level of cord (right) from an RO injected HCT47 animal. Inset above panel F shows high magnification view of intact DRGs still attached to the spinal cord. Tissue autofluorescence is observable in red channel. (**G-H**) Quantification of percent (%) specificity and completeness of labeling by the HCT47 virus in cervical spinal cord. Data are plotted as mean ± S.E.M and n = 4 animals and 3 sections per animal per level of cord were analyzed. (**I**) Representative Western blot from HCT1 and HCT47 RO injected animals showing SYFP2 protein levels in cervical spinal cord tissue lysates, revealed using an anti-GFP antibody, and β-actin protein levels as loading control for each sample. Each lane represents SYFP2 and β-actin from a single animal; n = 3 animals represented per virus. (**J**) Quantification of HCT47-driven SYFP2 protein levels relative to HCT1-driven levels. Data are presented as mean ± S.E.M for n= 5 animals per viral group. *p<0.05 for comparison of HCT1 vs HCT47 by unpaired two-tailed Student’s t-test.

We performed a sequence-based analysis of the 1137m enhancer to identify putative transcription factor (TF) binding motifs. This analysis revealed multiple significant TF binding domains, including Isl2 (p-value = 1.79e-05) which is a known regulator of MN development supporting the MN specific expression of 1137m and may be useful for future rationale *in silico* design of synthetic enhancer sequences (**Table S2**).

### Generation of enhancer-driven fluorophore AAVs for LMN subtypes

Somatic LMNs can be further divided into alpha, gamma and beta types depending on the type of muscle fiber innervated **(Figure 6A**)^6^ and have been shown to be differentially susceptible in various spinal cord diseases^9,28,29^. Therefore, we next sought to develop enhancer AAVs for alpha and gamma LMN subtypes for more refined targeting approaches. We discovered differentially accessible putative enhancers proximal to known marker genes, *Chodl* and *Ret,* for each subtype within the mouse and NHP spinal cord multiome datasets (**Figure 6B**) and proximal to other previously described marker genes (**Figure S3B)**^1,2,30^. We cloned 20 putative alpha LMN subtype and 3 putative gamma LMN subtype enhancers into the SYFP2-expressing rAAV backbone and screened PHP.eB viruses in wild-type mice following RO injection (**Figure 6C, D** and **Figure S3A, B**). We found 16 out of 20 alpha subtype LMN enhancers (80%) showed SYFP2+ labeling of very large cell bodies within transverse sections from cervical spinal cord, and out these, 10 labeled cells within the ventral horn (50%), which is consistent with the described location and morphology of alpha motor neurons (**Figure 6D and Figure S3A, B**). For putative gamma subtype LMN enhancers, we found that 3 out of the 3 (100%) showed SYFP2+ labeling of very small cell bodies within the ventral horn in transverse cervical cord sections, which is consistent with the described morphology (**Figure 6D and Figure S3A, B**). The HCT46-1188m-SYFP2 virus (referred to as HCT46 from here forward) exhibited the strongest and most abundant labeling of potential gamma subtype LMNs and therefore this virus was selected for further characterization. The HCT3-1139m-SYFP2 virus initially exhibited very specific labeling of potential alpha subtype LMNs but appeared somewhat weak. Therefore, we optimized it by concatenation of the putative enhancer core as previously described^16^, and created a derivative virus termed HCT49-3Xcore-1139m-SYFP2 (referred to as HCT49 from here forward; **Figure 6C**) for further characterization.

**Figure 6.**
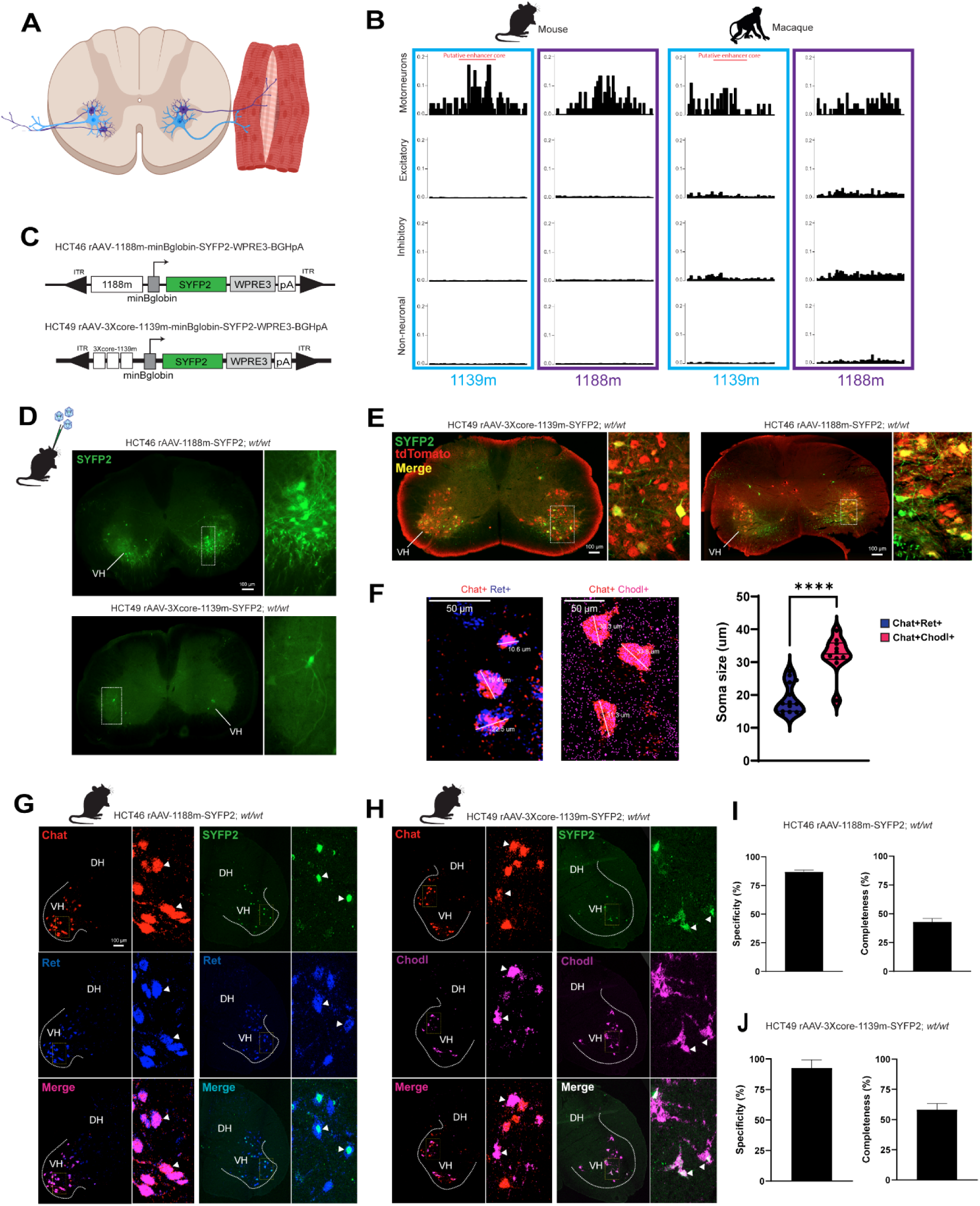
Generation and validation of enhancer AAVs for selective labeling of alpha or gamma spinal motor neurons in mouse cord. (**A**) Schematic of transverse cervical spinal cord section with alpha motor neurons (blue) and gamma motor neurons (purple) residing in ventral horn and sending projections to extrafusal or intrafusal skeletal muscle fibers, respectively. (**B**) Chromatin accessibility for 1139m and 1188m putative enhancers in mouse and macaque spinal cord. Each individual tract represents the indicated spinal cord cell type, and the y-axis is normalized accessibility within the genomic region of each enhancer. (**C**) HCT46 and HCT49 vector diagrams. The 1188m enhancer or a 3Xcore of the 1139m enhancer was cloned into the standard rAAV backbone. (**D**) Representative epifluorescence images of RO-injected, HCT46- and HCT49-driven native SYFP2 fluorescence in 50µm thick transverse spinal cord sections, respectively. VH = ventral horn and dashed box denotes area of interest highlighted in high magnification images to the right. (**E**) Representative epifluorescence images of native SYFP2 (green) signal and anti-Chat signal (red) in 30µm thick transverse cervical spinal cord sections in mouse. Inset panels to the right show cells boxed in low magnification image. (**F**) Representative confocal RNAscope images of Chat+/Ret+ gamma motor neurons (red/blue) labelled by the HCT46 virus and Chat+/Chodl+ alpha motor neurons (magenta/red) labelled by the HCT49 virus (left). White lines across the representative cells denote the calculated soma diameter. Average soma diameters for Chat+/Ret+ cells (2 mice, 7-8 sections each) and Chat+Chodl+ cells (4 mice, 2-4 sections each) were measured and are presented as the mean ± S.E.M ****p<0.0001, Mann Whitney test (**G-H**) Representative confocal RNAscope images of hemisected transverse cervical cord sections of 20µm thickness from HCT46 (**G**) and HCT49 (**H**) virus injected animals. Probes for *Chat* (red), *Ret* (blue), *SYFP2* (green) and *Chodl (magenta)* were used in two-color experiments to confirm probe specificity and validate each virus. Curved dashed lines indicate the ventral horn (VH) and DH = dorsal horn; Dashed yellow box in hemisected images denote the region of interest in the panels to the right. White arrowheads indicate some of the double positive cells (e.g., *Chat+*/*Ret+*) for each condition. (**I-J**) Quantification of HCT46 (**I**) or HCT49 (**J**) virus specificity and completeness of labeling in cervical mouse spinal cord sections. Data are plotted as mean ± S.E.M and n = 2 animals per virus and 3-8 sections per animal were analyzed.

To validate HCT46 and HCT49 viruses, we evaluated native SYFP2 fluorescence and anti-Chat signal co-localization in wild-type and in Chat-IRES-Cre;Ai75 double transgenic mice following RO injection of each virus. Here we found strong co-localization of SYFP2+ cells with Chat+ cells in the ventral horn of transverse cervical spinal cord sections, confirming labeling of a subset of LMNs (**Figure 6E and Figure S6F, G**). To validate each virus at the level of subtype, we performed RNAscope analysis on fresh frozen cervical spinal cord sections using a combination of *SYFP2*, *Chat*, and *Ret* or *Chodl* probes for the HCT46 or the HCT49 viruses, respectively (**see Methods Details**; **Figure 6G, H**). Here we observed the expected co-labeling of *Chat* signal with *Ret* or *Chodl* signals in single cells within HCT46 virus-infected sections (**Figure 5G, left panels**) or HCT49-virus infected sections (**Figure 5H, left panels**), which confirmed subtype probe specificity. Additionally, analysis of the soma diameter size of *Chat+/Ret+* or *Chat+/Chodl+* neurons revealed an average diameter of 18.6 ± 1.4 µm for gamma type MNs and 32.1 ± 1.7 µm for alpha type MNs, which is generally consistent with those reported^31^ (**Figure 6F**). We then evaluated the specificity of the HCT46 or HCT49 viruses and found a high degree of overlap between *SYFP2+* and *Ret+* or *Chodl+* cells in cervical spinal cord sections (**Figure 6G, H, right panels**). Analysis of these data revealed that both HCT46 and HCT49 viruses exhibit high specificity for gamma and alpha subtype LMNs (86.9% and 92.5%, respectively; **Figure 6I, J, left panels**). Additionally, we found that more than half of alpha subtype LMNs were labeled by the HCT49 virus (58.1%; **Figure 6J, right panel**) and 42.9% of gamma subtype LMNs were labeled by the HCT46 virus in cervical spinal cord sections (**Figure 6I, right panel**).

### Enhancer stitching enables dual labeling of UMNs and LMNs by a single virus

We have previously shown that enhancer viruses targeting different cell types can be co-delivered to achieve simultaneous labeling in one animal^16^. To extend this approach and simplify it, we next sought to develop a single vector system to achieve combinatorial labeling of cell types within the brain and spinal cord. We targeted L5 ET neurons of the corticospinal tract with LMNs (**Figure 7A**) because these cell types selectively degenerate in diseases such as ALS)^28,32^, suggesting potential therapeutic applications for such a vector. This approach was also pursued because we have a large collection of well validated enhancers for L5 ET neurons^16^ and LMNs (present study). We cloned the L5 ET neuron enhancer 453m upstream of the LMN 1137m enhancer and the minimal beta-globin promoter in a SYFP2-expressing viral vector (**Figure 7B**). We then packaged HCT55-453m-1137m-SYFP2 (referred to as HCT55 from here on) and 453m or 1137m single enhancer control vectors into PHP.eB serotyped viruses and performed RO injections in wild-type mice (**Figure 7C**). As expected, the 453m enhancer alone drove robust native SYFP2 fluorescence of L5 ET neurons within cortex throughout the full rostral caudal axis, albeit strongest in posterior cortex, and in neurons within the deep cerebellar nucleus (DCN) and within lamina 10 (10Sp) of the spinal cord (**Figure 7C, left panels**). The 1137m enhancer alone drove robust SYFP2 expression in LMNs and in several hindbrain structures (**Figure 7C, middle panels**), as predicted. With the HCT55 stitched virus, we found the combined robust native SYFP2 expression in L5 ET neurons throughout the cortex and in neurons enriched in the ventral horn of the spinal cord (**Figure 7C, right panels**). Many of these ventral horn neurons were confirmed to be LMNs following additional validation using anti-Chat immunostaining or Chat-IRES-Cre;Ai75 double transgenic mice (**Figure 7D and Figure S6H**). Quantitative analysis of SYFP2+ and Chat signal co-localization revealed highly specific viral labeling in MMC (86.4-93.2%), LMC (77.7-89.0%), Ph9 (100%) and completeness of labeling ranging from 60.6-95.0% in the same areas (**Figure 7I, J**). Sparse and incomplete viral labeling of PCs and in PGC was observed in at least one level of spinal cord or a total absence of viral labeling (**Figure 7I, J**). With more coarse segmentation, quantitative analyses revealed that the HCT55 virus labeled 40.8% of LMNs in cervical cord with 60.3% specificity (**Figure S6I, J**), which likely only reflects the lower resolution anatomical analysis method (**Figure S6I, J**).

**Figure 7.**
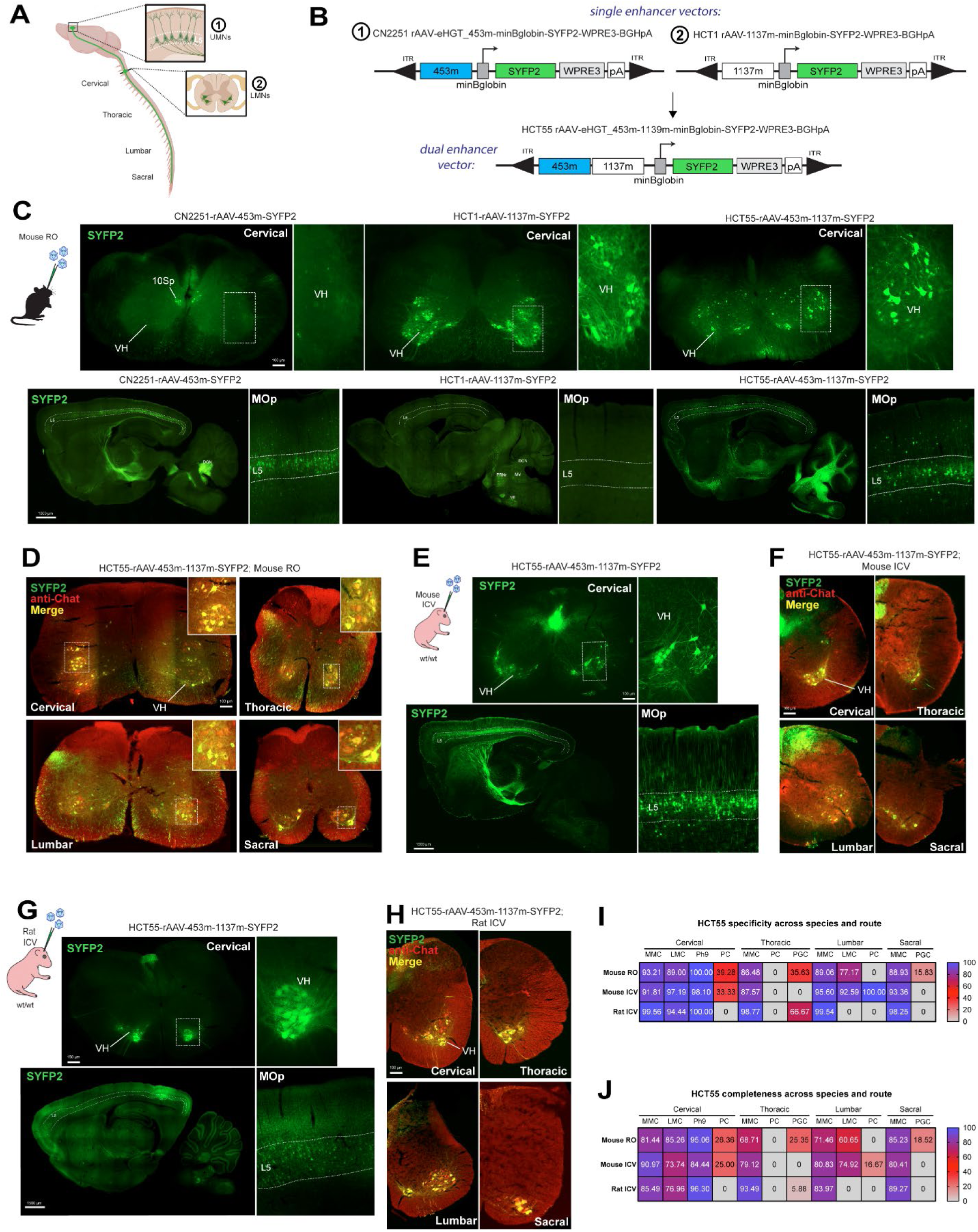
Enhancer stitching enables simultaneous labeling of L5 ET neurons and spinal motor neurons with a single viral vector. (**A**) Schematic of mouse central nervous system showing brain and spinal cord. L5 ET neurons located within motor cortex (M1) are highlighted in the 3D brain structure and the right panel (1), and Lower motor neurons (LMNs) located in the ventral horn of the spinal cord are highlighted in a transverse cervical section in the right panel (2). The line traveling from brain to spinal cord represents the corticospinal tract. (**B**) Schematics of CN2251, HCT1, and HCT55 vector designs. In the single enhancer vectors (CN2251 and HCT1), the 453m and 1137m enhancer sequences were cloned into the standard backbone upstream of the minBglobin promoter. In the dual enhancer vector (HCT55), the 453m enhancer was cloned directly upstream of the 1137m enhancer. (**C**) Representative epifluorescence images of virus driven SYFP2 expression in 50µm thick transverse spinal cord sections (upper panels) and in 25µm thick sagittal brain sections (lower panels). VH = ventral horn, 10Sp = lamina 10, DCN = deep cerebellar nucleus, PRNr = pontine reticular nucleus, MV = medial vestibular nucleus, VII facial motor nucleus, and the dashed boxes indicate the region of interest highlighted in the right panels. Dashed lines on the sagittal brain sections denote cortical L5 across the full rostral-caudal axis and representative images from M1 are shown in the right panels. (**D**) Representative epifluorescence images of native SYFP2 fluorescence (green) and anti-Chat staining (red) in 30µm thick transverse spinal cord sections at every level of cord from HCT55 virus RO-injected mice. (**E**) Representative epifluorescence images of SYFP2 fluorescence in 50µm thick transverse cervical spinal cord sections or 25µm thick brain sagittal sections following ICV delivery of the HCT55 virus into mouse neonatal pups. (**F**) Representative epifluorescence images of native SYFP2 (green) fluorescence and anti-Chat signal (red) in 30µm thick transverse spinal cord hemi-sections at every level of cord from HCT55 virus ICV-injected mice. (**G**) Representative epifluorescence images of SYFP2 fluorescence in 30µm thick transverse cervical spinal cord sections or 25µm thick brain sagittal sections following ICV delivery of the HCT55 virus into rat neonatal pups. n = 2 animals and 3 sections per animal analyzed. (**H**) Representative epifluorescence images of native SYFP2 (green) fluorescence and anti-Chat signal (red) in 30µm thick transverse spinal cord hemi-sections at the indicated level of cord from HCT55 virus ICV-inject neonatal rat pups. (**I-J**) Heatmap representation of percent (%) specificity and completeness of HCT55 viral labeling of Chat+ motor neurons in the spinal cord following RO injection in mouse or ICV injection in mouse or rat pups. Legend with scale denotes values ranging from 0 (grey) – 100% (purple). Percentage values for all mouse experiments are the average mean value from n = 3 animals and 3 sections per animal per level of cord analyzed. Percentage values for all rat experiments are the average mean value from n=1 animal and 3 sections per animal per level of cord analyzed.

We additionally tested the HCT55 virus via the ICV route of administration in mouse neonatal pups and evaluated native SYFP2 fluorescence in the brain and spinal cord several weeks post-injection (**Figure 7E, G**). Here we found extremely robust labeling of L5 ET neurons throughout the cortex and a notable decrease in the amount of L2/3 pyramidal neuron expression in primary motor cortex (MOp) compared to that achieved with mouse RO injection (**Figure 7E, G bottom panels**). Similarly to RO condition, we observed highly selective labeling of LMNs within MMC (87.5-98.1%), LMC (92.5-97.1%), Ph9 (98.1%), and in PCs (100% at lumbar level) (**Figure 7I**). Completeness of labeling ranged from 73.7-90.9% in MMC, LMC and Ph9 regions and from 0-25% in PCs and PGC (**Figure 7J**).

Lastly, we evaluated the ability of the HCT55 virus to label both L5 ET neurons and LMNs across rodent species. The virus was injected into rat neonatal pups via the ICV route of administration and native SYFP2 fluorescence was evaluated in the target tissues several weeks post-injection (**Figure 7G**). In these experiments, we observed robust and specific SYFP2 labeling of LMNs in cervical spinal cord (**Figure 7G, top panels**) and labeling of L5 ET neurons with most prominent expression within caudal regions of the brain (**Figure 7G, bottom panels**). Anti-Chat immunostaining was performed and robust co-localization of SYFP2 fluorescence with Chat signal was observed throughout all levels of spinal cord (**Figure 7H**). Quantification of these data revealed highly specific labeling in MMC (98.2-99.5%), LMC (94.4% cervical level), and Ph9 (100%) (**Figure 7J**). Completeness of labeling ranged from 76.6-96.3% in MMC, LMC, and Ph9 regions and from 0-5.8% in PCs and PGC (**Figure 7J**). Taken together, these data demonstrate that the individual enhancer activity is conserved between rodents in the stitched vector design and largely similar results can be obtained with different routes of administration.

### Generation of LMN enhancer AAVs expressing iCre297T or ChR2(H134R)-EYFP

We have previously shown that enhancer AAVs expressing recombinases (e.g. Cre, Flp, Nigri) can be used with existing transgenic reporter lines or viral reporters to access a diversity of brain cell types^16^. To extend this toolkit beyond the brain, we next developed pan- or subtype-selective LMN viruses expressing an attenuated iCre (iCre297T; HCT157, HCT158 and HCT160) and tested them by RO injection of Ai14 Cre-dependent tdTomato reporter animals (**Figure 8A**). Each virus drove the expected selective recombination pattern in Chat+ neurons located within the ventral horn at the level of cervical spinal cord (**Figure 8B**). Quantitative analysis of HCT157 driven tdTomato fluorescence and anti-Chat signal co-localization in cervical cord revealed high levels of specificity and completeness across MMC (89.7 and 87.8%), LMC (89.7 and 82.1%), and Ph9 (97.6 and 90.3%) (**Figure 8C, D**). Like HCT1, specificity and completeness of HCT157 viral labeling of PCs was much lower than in the other areas analyzed (16.6 and 16.6%) (**Figure 8C, D**). HCT158 and HCT160 viruses labeled subsets of Chat+ neurons with a soma size of 18.6 ± 1.1 µm or 26.6 ± 0.6 µm, suggesting effective targeting of gamma and alpha type MNs, respectively (**Figure 8E**).

**Figure 8.**
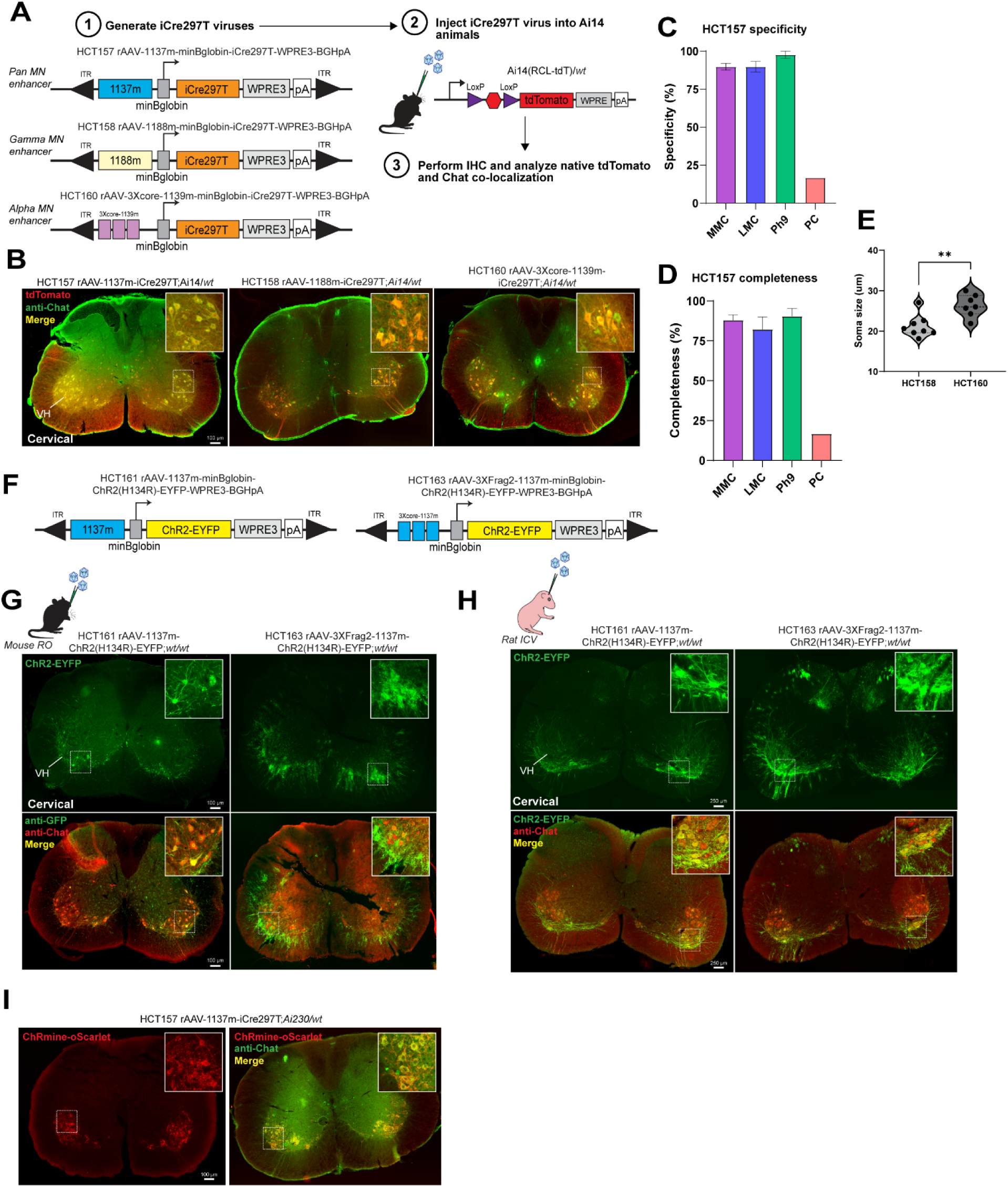
Generation and validation of Cre- and Opsin-expressing enhancer AAVs for selective targeting of spinal motor neurons. (**A**) iCre297T enhancer vector designs and experimental workflow. An iCre297T gene was cloned in place of the SYFP2 gene in MN vectors HCT157, HCT158 and HCT160. PHP.eB serotyped viruses were generated and injected into Ai14 reporter mice and native tdTomato fluorescence was analyzed. (**B**) Representative images of native tdTomato fluorescence (red) and anti-Chat signal (green) in 30µm thick transverse cervical spinal cord sections from Ai14 heterozygous reporter mice injected RO with the indicated virus. VH = ventral horn and dashed box denotes area of interest highlighted in high magnification images to the right. (**C**) Quantification of percent (%) specificity and completeness of HCT157 viral labeling of Chat+ motor neurons in cervical cord in anatomical areas and previously defined cell types following RO injection. MMC = medial motor column, LMC = lateral motor column, Ph9 = phrenic motor neurons of lamina 9, and PC = partition cells. Data are plotted as mean ± S.E.M and n = 3 per virus and 3 sections per animal were analyzed. (**E**) Average soma diameters for cells labelled by HCT158 or HCT160 viruses. Data are presented as the mean ± S.E.M; n = 3 animals per virus and 2-3 sections per animal were analyzed (**p=0.0059, Mann Whitney test). (**F**) Schematics for ChR2(H134R)-EYFP2 expressing vectors. A ChR2(H134R)-EYFP2 was cloned in place of the SYFP2 gene in the 1137m and optimized 3Xcore-1137m enhancer vectors. PHP.eB serotyped viruses were generated and injected RO into wild-type mice or ICV into neonatal rat pup. (**G-H**) Representative epifluorescence images of native SYFP2 alone (green in upper panels) or native SYFP2 and anti-Chat signal (red in lower panels) in 30µm thick transverse cervical spinal cord sections from RO injected mice (**G**) or ICV injected rats (**H**). VH = ventral horn and dashed box denotes area of interest highlighted in high magnification images to the right. (**I**) Representative epifluorescence images of native soma enriched ChRmine-oScarlet fluorescence (red in left panel) or soma enriched ChRmine-oScarlet and anti-Chat signal (green in right panel) in 30µm thick transverse cervical spinal cord sections from Ai230 heterozygous mice injected RO with the HCT157 virus. High magnification inset are of the boxed ventral horn cells in the low magnification image.

To enable functional experiments, we lastly generated enhancer AAVs for pan LMN targeting that express channelrhodopsin-2 (ChR2) with H134R mutation fused to EYFP (ChR2(H134R)-EYFP) (**Figure 8G, H**). 1137m- and 3XFrag2-containing enhancer vectors (HCT161 and HCT163, respectively) were generated and tested by RO injection into wild-type mice (**Figure 8G**) or ICV injection into rat neonates (**Figure 8H**). We found that both viruses drove robust native expression of ChR2(H134R)-EYFP fluorescence in the ventral horn of cervical spinal cord sections in both species (**Figure 8G, H, upper panels**), and virally labelled cells were Chat+ LMNs (**Figure 8G, H, bottom panels**). 1188m- and 3XCore-1139m-enhancer AAVs driving ChR2(H134R)-EYFP were also generated and tested in parallel, but only weak and sparse native fluorescence was observed for both suggesting additional vector optimization is needed to achieve functionally relevant transgene levels (data not shown). As a complimentary approach, we also evaluated whether the HCT157 virus could be used in conjunction with the Ai230 Cre-dependent soma-enriched ChRmine-oScarlet transgenic line^33^ to drive opsin expression selectively in LMNs (**Figure 8I**). Ai230 heterozygous animals were RO injected with HCT157 and native ChRmine-oScarlet expression in spinal cord was evaluated, which revealed strong, native ChRmine-oScarlet expression predominantly within the soma of Chat+ LMNs in cervical spinal cord (**Figure 8I**). Taken together, these data demonstrate that the enhancers validated for pan- and subtype LMNs can be utilized to drive a variety of different cargoes into their respective target neuronal populations.

## Discussion

Currently available single-cell ATAC-seq data sets for analyzing chromatin accessibility in the rodent spinal cord are limited to very early in development^34–36^ and are non-existent in the case of primate spinal cord. Therefore, the 10X multiome datasets, which reveal both the transcriptome and epigenome in the same cell, generated in the present study from adult mouse and adolescent macaque represent a unique and tremendously valuable resource to the community. We have provided a roadmap for how these datasets can be used to identify functional enhancers and created a suite of MN selective viral tools for labeling of cell types in the adult rodent and NHP spinal cord. With additional analyses, these multiome datasets may enable enhancer discovery and tool development beyond MNs. This has the potential to re-define genetic approaches for targeting a diversity of spinal cord cell types and overall cell type definition across species.

Our previous efforts established the enhancer AAV technology platform as a robust approach for brain cell type-specific AAV tool development^15–18^. In the present work, we used similar methodology for one at a time screening and validation of candidate enhancers for pan MNs and MN subtypes. We had a high success rate for translating enhancer discovery into functional tools (> 75%), with equivalent or greater than we had previously targeting brain cell types. These results, combined with our success developing enhancer AAVs for targeting cell types outside of the CNS (data not shown), suggest that any cell type within the body is within reach. Future work in this space will rely heavily on the availability of high quality epigenomic data with sufficient cell type resolution such as previously reported^21,37^ and optimization of other experimental parameters (i.e. capsid, delivery route, and dose) to ensure success.

The single 1137m enhancer-containing viruses (HCT1 and HCT69) provided highly specific labeling of LMNs located in MMC, LMC, and Ph9 in mouse, rat, and macaque spinal cord following different routes of administration. Variations in the extent of labeling (i.e., completeness) were minimal between RO and ICV routes of administration in mouse and between ICV injected mouse versus rat. Similarly to in rodents, both ICM and IT routes of administration in macaque yielded highly specific viral labeling in the same anatomically defined regions. The completeness of labeling with both routes of administration was more moderate in macaque which likely reflects the dose of virus tested, although we cannot fully confirm that without an additional replicate for each condition. We think it’s likely that the completeness of LMN labeling in macaque spinal cord could be improved with delivery of a higher dose of virus by either route of administration. Nonetheless, we are not aware of any other viral tools that can be used singly to label LMNs to this degree in the adult mouse, let alone across three mammalian species and therefore, these tools are incredibly unique and likely to be enabling for direct analysis of LMN function and connectivity in subsequent studies.

For potential translational applications, such as for human gene therapy, we evaluated 1137m enhancer-driven expression in many peripheral organs of the mouse and macaque and found essentially no expression outside of LMNs within the spinal cord and this was notably more specific than that observed with a hSyn1 promoter-driven transgene. A lack of off-target viral expression in DRGs and vital organs such as the heart and liver are desirable for potential gene therapies, as it is expression there with high doses of CAG- or hSyn1-promoter driven viruses that often underlie the clinical failures^38^. Therefore, the 1137m enhancer has the ideal body-wide biodistribution profile, which suggests that it could be deployed in therapeutic strategies and delivered less invasively as a PHP.eB serotyped virus via ICM or IT routes to access LMNs throughout the entire primate spinal cord.

Differential vulnerability of LMN subtypes in neurodegenerative diseases is a recurring theme and therefore, we generated enhancer AAVs targeting alpha or gamma type LMNs and showed high selectively of these viruses for *Chodl*+ fast-firing (FF) alpha type LMNs and *Ret+* gamma type LMNs respectively in mouse spinal cord. Much less is known about *Chodl*+ FF alpha type LMNs and *Ret*+ gamma subtype LMNs and other transcriptomically-defined LMN subtypes due to their recent discovery^2^ and the lack of available genetic tools to selectively access these cell types. It is generally accepted that at the broader subclass level alpha type LMNs selectively degenerate in diseases such as ALS, while gamma type LMNs are spared^8,39^. Therefore, subtype-selective LMN interventions enabled by our enhancer AAVs may be beneficial to slow disease progression and/or to provide support to the spared neuronal population. Accessibility of the functionally characterized subtype selective LMN enhancers were found to be conserved in the macaque spinal cord multiome dataset and therefore they may be portable for use in primate.

The ability to target multiple cell types selectively and simultaneously in a flexible manner using a single enhancer-based viral vector offers many advantages and increases the potential for use in therapeutic approaches. As proof of concept for this general approach, and for use in the ALS disease context, we stitched validated enhancers for L5 ET neurons and LMNs together to target descending motor pathways and found that the combined labeling pattern could be achieved in the CNS. The specificity and completeness of viral labeling of LMNs located in MMC, LMC, and Ph9 was very high at most spinal cord levels evaluated following RO or ICV delivery to mice or ICV delivery to rats. Some subtle differences in viral labeling patterns in the cortex between routes of administration in mice and between rodent species were observed, such as reduced layer 2/3 excitatory neuron labeling with mouse ICV injection and presence of layer 6b excitatory neuron labeling with rat ICV injection. Additionally, we consistently observed qualitatively higher levels of reporter expression following ICV injection to rodents. Collectively these observations likely reflect differences in the multiplicity of infection (MOI) obtained on a single cell basis with the different routes of administration but may also suggest differences in the underlying gene regulatory network or capsid tropism. Given that both the individual L5 ET neuron and LMN enhancers contained within the stitched design have been shown to label the homologous cell type in primate spinal cord and brain (as demonstrated in the current paper and additional data not shown), it seems likely that the combined enhancer vector would exhibit the same specificity. This combined enhancer vector could then be used with a recently discovered BBB-penetrant capsid (e.g. bCap1, Dyno Therapeutics unpublished) to achieve simultaneous targeting of a therapeutic cargo with a less invasive intravenous (IV) route of administration.

For functional studies, we created a toolkit of viruses expressing iCre297T or ChR2(H134R)-EYFP in LMNs and demonstrate that these can be used with existing mouse transgenic lines for cell type labeling or in principle for *in vitro* or *in vivo* physiology or behavioral experiments. The ChR2(H134R)-EYFP viruses performed as well in rats as in mouse and could likely be used directly for similar cell- and circuit-level investigations in these rodent species or potentially in other mammalian species.

The large suite of enhancer AAVs reported here are first-in-class for the spinal cord field and will enable diverse cell type-specific and systems-level applications, which will ultimately lead to a better understanding of MN function in both healthy and diseased states, and potentially more effective therapeutic interventions for debilitating and lethal neurodegenerative disorders.

## Supporting information

Key Resource Table

## Acknowledgements

We are grateful for the support of the following Allen Institute teams and departments: Transgenic Colony Management, Genotyping, Lab Animal Services, Viral Technology Core, Tissue Processing, Molecular Biology, Technology, and Human Cell Types. We thank Leah Chestnut and Brian Nguyen at LifeCanvas Technology for light sheet imaging, video, and image preparation. We thank members of the Washington National Primate Research Center for providing animal care, surgical support, and assistance with tissue collection for the study. We thank Jim Berg for providing mouse intraspinal injection methods. Schematics in many of the figures were created with BioRender.com. This work was supported by award number U19MH114830 from the National Institute of Mental Health (NIMH) to H.Z. and by BioMarin Pharmaceutical and the Paul G. Allen Foundation. The WaNPRC is supported by the National Institutes of Health (NIH), Office of Research Infrastructure Programs (ORIP) under award number P510D010425 *and U420D011123.* Its contents are solely the responsibility of the authors and do not necessarily represent the official views of the NIH, the NIMH or of BioMarin Pharmaceutical. We thank the Allen Institute founder, Paul G. Allen, for his vision, encouragement, and support. This publication was supported by and coordinated through the <u>BRAIN Initiative Armamentarium for Precision Brain Cell Access</u> consortium.

## Author contributions

T.L.D designed the study. R.H., C.S., and R.F. performed macaque spinal cord tissue collections, R.H., T.E.B., and N.J. oversaw 10X multiome generation from macaque, and performed analyses. N.D. managed tissue processing team and T.C., M.C., W.H., performed mouse spinal cord tissue collections for multiome data generation. Z.Y., Y.G., C.T.J.vV., B.T., N.J., and H.Z. oversaw 10X multiome generation from mouse and performed analyses. E.K, N.T., A.C., A.F., T.L.D., B.W., R.M., E.L.G., performed molecular cloning, histology, imaging, and analysis. N.T. M.R.B, B.C.C., C.E., J.C.D., and M.G.F. performed macaque injections and collected and processed macaque tissue. S.Y., N.D., and D.N. generated viruses. M.R., E.L, L.S., and C.H., performed mouse tissue collections. M.A.Q., A.G.C., E.K. and T.L.D. performed western experiments and analyses. E.K. performed RNAscope experiments and analyses. A.C. performed soma size analyses. J.T.T. oversaw rat experiments and protocol, and with E.K. performed tissue collection, imaging, and analyses. J.T.T. and K.G. managed macaque experiments and protocol. C.R., A.D. and K.D. collaborated with T.L.D., H.Z., and B.T. to generate the Ai230 transgenic mouse line. A.W. managed the neurosurgery and behavior team members R.N. and J.K. that performed intraspinal injections. K.A.S. managed molecular biology team members A.B.C., J.G., A.T., J.B.G., R.C., B.N., N.G., T.H.P. and multiome data production. E.S.L. provided leadership of the Human Cell Types department and the CNS Gene Therapy program. J.T.T., B.P.L., T.L.D. provided leadership of the CNS Gene Therapy and Viral Genetic Tools programs. T.L.D. wrote the manuscript with support from many co-authors.

## STAR Methods

### Resource availability

#### Lead Contact

Further information and requests for resources and reagents should be directed to and will be fulfilled by the Lead Contact, Tanya L. Daigle.

#### Materials availability

All plasmids generated in this study have been deposited to Addgene.

#### Data and Code availability

Newly generated 10x Multiome datasets from mouse and macaque have deposited to NeMO. The Nemo identifier is nemo:dat-vccmgkw and can be found at https://assets.nemoarchive.org/dat-vccmgkw. Software code used for data analysis and visualization is available from GitHub at https://github.com/AllenInstitute/kussick2024analysis/. R packages for constructing and annotating cell type taxonomies can be found on GitHub at https://github.com/AllenInstitute/scrattch.taxonomy and https://github.com/AllenInstitute/scrattch.mapping respectively.

### Experimental model and subject details

#### Mouse breeding and husbandry

Adult C57Bl/6J mice were bred and housed under Institutional Care and Use Committee (IACUC) protocols 2202 and 2301 at the Allen Institute for Brain Science, with no more than five animals per cage, maintained on a 12 day/night cycle, with food and water provided *ad libitum*. Juvenile C57BL/6J mice were also purchased directly from The Jackson Laboratory (Strain #000664) and housed at the Allen Institute for Brain Science. Both male and female mice were used for experiments and the minimal number of animals was used for each experimental group. Animals with anophthalmia or microphthalmia or other obvious conditions (e.g. penile prolapse) were excluded from experiments and all animals were maintained on a C57BL/6J background.

The Chat-IRES-Cre knock-in mouse line was purchased from The Jackson Laboratory (Strain #006410) and maintained on a C57BL/6J background. The *Rosa26* reporter line Ai75 (RCL-nls-tdTomato) line was generated in house^40^ and maintained on a C57BL/6J background. Only mice heterozygous for both reporter and Cre driver were used for experiments. Ai230 TIGRE transgenic reporter mice (TIT2L-XCaMPG-WPRE-ICL-ChRmine-TS-oScarlet-Kv2.1-ER-IRES2-tTA2-WPRE_hyg strain #37944) were generated in-house^33^ under IACUC protocol 2453 and maintained on a C57BL/6J background.

#### Rat husbandry

Timed-pregnant female CD-1 Sprague-Dawley rats were purchased from Charles River Laboratories (Strain code #001) and utilized for experiments under Allen Institute Institutional Care and Use Committee protocol #2010. Animals were housed no more than 1 per cage, maintained on a 12 day/night cycle, with food and water provided *ad libitum*. Rat pups were born generally the week following shipment receipt and at P1, the pups were tattooed for identification purposes and used for unilateral ICV injection of AAV vectors.

#### Macaque husbandry

Surgical procedures, experimental protocols and animal care conformed to the NIH Guide for the Care and Use of Laboratory Animals and were approved by the Institutional Animal Care and Use Committee at the University of Washington. IACUC Protocol Number 4532-10. Animal husbandry and housing were overseen by the Washington National Primate Research Center. Monkeys had *ad libitum* access to biscuits (Lab Diet, Fiber Plus Monkey Diet Cat#5049) and controlled daily access to fresh produce, treats and water. When possible, animals were pair-housed and allowed grooming contact. Cages were washed every other week, bedding was changed every day, and animals were examined by a veterinarian at least twice per year.

### Methods details

#### 10X multiome data generation from mouse and macaque spinal cord

Macaque tissue samples were obtained from the University of Washington National Primate Resource Center. Immediately following euthanasia, macaque spinal cord samples were removed and transported to the Allen Institute in artificial cerebral spinal fluid equilibrated with 95% O_2_ and 5% CO_2_. Upon arrival at the Allen Institute, spinal cord samples were flash frozen in dry-ice cooled isopentane, transferred to vacuum-sealed bags, and stored at -80°C. To isolate specific regions of interest, spinal cord samples were briefly transferred to -20°C and the region of interest was removed and subdivided into smaller blocks on a custom temperature controlled cold table held at -20°C. After dissection, tissue blocks were returned to -80°C until the time of Multiome processing. Nucleus isolation for 10x Chromium Multiome was conducted as described (dx.doi.org/10.17504/protocols.io.y6rfzd6). Briefly, single nucleus suspensions were incubated with DAPI (4’,6-diamidino-2-phenylindole dihydrochloride, ThermoFisher Scientific, D1306) at a concentration of 0.1µg/ml, Alexa Fluor 488 rabbit anti-OLIG2 antibody (Abcam ab225099, 1:2000), and mouse anti-NeuN conjugated to PE (FCMAB317PE, EMD Millipore. 1:500). Controls were incubated with mouse IgG1k-PE Isotype control (BD Biosciences, 555749, 1:250 dilution), Alexa Fluor 488 rabbit IgG Isotype Control (Abcam ab199091, 1:250) or DAPI alone. Single-nucleus sorting was carried out on either a BD FACSAria II SORP or BD FACSAria Fusion instrument (BD Biosciences) using a 130 µm nozzle and BD Diva software v8.0. A standard gating strategy based on DAPI, NeuN, and OLIG2 staining was applied to all samples. Doublet discrimination gates were used to exclude nuclei multiplets. NeuN+, OLIG2+, and OLIG2-nuclei were sorted into separate tubes and were pooled at a targeted ratio of 70% NeuN+, 20% OLIG2-, and 10% OLIG2+ after sorting. Sorted samples were centrifuged and the concentration of nuclei (nuclei/µl) was determined. Nuclei were immediately loaded for 10x Chromium multiome processing following the manufacturer’s protocol.

Mice were anaesthetized with 2.5–3% isoflurane and transcardially perfused with cold, pH 7.4 HEPES buffer containing 110 mM NaCl, 10 mM HEPES, 25 mM glucose, 75 mM sucrose, 7.5 mM MgCl2, and 2.5 mM KCl to remove blood (see *Protocols.io* https://doi.org/10.17504/protocols.io.5jyl8peq8g2w/v1). Following perfusion, the spinal cord was dissected quickly, cut into 4 parts (C1-C8, T1-T13, L1-L6, S1-S4 + Co1-Co3), frozen for 2 min in liquid nitrogen vapor and then moved to −80 °C for long term storage following a freezing protocol developed at AIBS (see *Protocols.io* https://doi.org/10.17504/protocols.io.j8nlkodr6v5r/v1).

Nuclei were isolated using the RAISINs method ^41^ with a few modifications as described in a nuclei isolation protocol developed at AIBS (see *Protocols.io* https://doi.org/10.17504/protocols.io.4r3l22n5pl1y/v1. In short, excised tissue dissectates were transferred to a 12-well plate containing CST extraction buffer. Mechanical dissociation was performed by chopping the dissectate using spring scissors in ice-cold CST buffer for 10 min. The entire volume of the well was then transferred to a 50-ml conical tube while passing through a 100-µm filter and the walls of the tube were washed using ST buffer. Next the suspension was gently transferred to a 15-ml conical tube and centrifuged in a swinging-bucket centrifuge for 5 min at 500 rcf and 4 °C. Following centrifugation, the majority of supernatant was discarded, pellets were resuspended in 100 µl 0.1× lysis buffer and incubated for 2 min on ice. Following addition of 1 ml wash buffer, samples were gently filtered using a 20-µm filter and centrifuged as before. After centrifugation most of the supernatant was discarded, pellets were resuspended in 10 µl chilled nuclei buffer and nuclei were counted to determine the concentration. Nuclei were diluted to a concentration targeting 5,000 nuclei per µl.

For 10x Multiome processing, we used the Chromium Next GEM Single Cell Multiome ATAC + Gene Expression Reagent Bundle (1000283, 10x Genomics). We followed the manufacturer’s instructions for transposition, nucleus capture, barcoding, reverse transcription, cDNA amplification and library construction (see *Protocols.io* https://doi.org/10.17504/protocols.io.bp2l61mqrvqe/v1).

#### Genome assemblies and annotations

*M.* musculus (mouse) assembly: mm10, GRCm39 annotation:GCF_000001635.27-RS_2023_04; *M. mulatta* (rhesus monkey) assembly: Mmul_10 (rheMac10), annotation: NCBI Annotation Release 103.

#### 10X Multiome RNA-seq data processing and analysis

Raw sequencing data were processed using cellranger-arc (10x Genomics) to generate single-nucleus RNA-seq (snRNA-seq) UMI count matrices for intronic and exonic reads. We annotated cell classes for mouse and macaque by clustering cells and finding enrichment of existing marker panels for broad spinal cord cell classes ^1^. Nuclei were filtered to remove low-quality samples by requiring ≥1000 genes detected per non-neuronal nuclei and requiring ≥2000 genes detected per neuronal nuclei. Counts were normalized to transcripts per million and log transformed using scanpy ^42^. Putative doublets were removed by setting a threshold of 0.3 on doublet score. Species correction to align mouse and macaque was performed using scVI. A k-nearest neighbour graph was built using the species corrected latent dimensions from scVI. To visualize cell classes, we performed the uniform manifold approximation and projections (UMAPs) nonlinear dimension reduction technique.

#### ATAC–seq peak calling and filtering

We used ArchR for snATAC–seq peak calling on pseudo bulk ATAC–seq fragments grouped by cell class per species using the standard ArchR processing pipeline. High quality nuclei were retained based on ATAC-seq metrics: transcription start site enrichment >= 3, fragments > 1000, max fragments < 10,000. Cell class annotations were transferred on the snATAC-seq from the snRNA-seq annotations based on distinct marker gene expression. Fragment files were aggregated into cell class bigwig files using ArchR and uploaded to the UCSC genome browser for identification of putative enhancer elements.

#### Accessibility screening in external data

Accessibility counter screening in human whole body and mouse whole body was performed by downloading or constructing bigwig files from ^21,22^. To assess the accessibility of each putative spinal cord enhancer in these previous studies we used ‘multiBigwigSummary’ from deeptools. This tool produces for each enhancer the average number of fragments within an enhancers genomic coordinate for each ATAC-seq study.

#### Viral vector construction

Putative enhancers were cloned from C57BL/6J mouse genomic DNA using enhancer-specific primers and Q5 high-fidelity DNA polymerase (NEB Cat#M0494S). Individual enhancer sequences were initially cloned into a rAAV backbone that contained a minimal beta-globin promoter, one of the three following inserts: SYFP2 (a yellow fluorescent gene), iCre297T (a cre recombinase with a R297T mutation gene^43^, or ChR2(H134R)-EYFP2 (a channel rhodopsin 2 with a H134R mutation directly fused to EYFP2 gene^44^, a bovine growth hormone polyA (BGHpA), and a woodchuck post-transcriptional regulatory element (WPRE3) using standard restriction digest cloning or Gibson assembly techniques. The 3xcore of the 1137m enhancer, the 1139m enhancer, and the 3XFlag-mTFP1 transgenes were synthesized by Genscript and inserted into the standard rAAV backbone using standard restriction digest cloning. All plasmid sequences were verified via Sanger sequencing and restriction digests were performed to confirm intact inverted terminal repeat (ITR) sites.

#### Viral packaging and titering

Crude and purified rAAVs of the PHP.eB, 9P31, or PHP.S serotypes were generated in house using previously described methods ^16^. The average titer of these viral preps was 5.0 × 10^13^ GC/ml.

#### Mouse retro-orbital (RO) and intracerebroventricular injections (ICV) of AAVs, histology, immunohistochemistry, and imaging

C57BL/6J, Ai14 heterozygous, Ai230 heterozygous, or Chat-IRES-Cre;Ai14 double transgenic mice were retro-orbitally injected in 21–45-day-old age range (P21-P45). Mice were briefly anesthetized with isoflurane and 5.0 × 10^11^ viral genome copies (GC) were delivered into the right venous sinus in a volume of 90µL or less. 0.5% proparacaine hydrochloride ophthalmic solution (Patterson Cat#07-885-9765) was then applied to the injected eye and animals recovered the same day. Mice were euthanized 3-4 weeks post-infection for analysis.

For ICV injections, C57BL/6J mouse neonatal pups (P0-P3) were injected with 1.0 × 10^11^ GC into the right cerebral ventricle and euthanized 3-4 weeks post-infection for analysis.

On the day of sacrifice, mice were anesthetized with isoflurane and transcardially perfused with 1x phosphate buffered saline (PBS) followed by 4% paraformaldehyde (PFA). Organs of interest were removed, post-fixed in 4% PFA overnight at 4°C, followed by an additional incubation for 2-3 days in 30% sucrose at 4°C. Transverse spinal cord sections or sagittal brain sections or varying angles for peripheral tissue sections were cut using a freezing microtome (Leica SM2000R) or cryostat (Leica CM3050S). Section thickness was 25-50µm across the various tissues and sections were stored in 1xPBS containing 0.01% sodium azide (Millipore Sigma Cat#S2002) at 4°C until downstream applications.

For SYFP2 and cholinergic positive neuron detection, fixed sections were washed three times in 1xPBS at room temperature with gentle shaking, blocked in 1xPBS containing 5% normal goat serum (Vector Laboratories Cat#S-1000-20), 0.2% Triton X-100 (Millipore Sigma Cat#X100), bovine serum albumin (Millipore Sigma Cat#A9418-5G) for up to 2 hours at room temperature with gentle shaking, and then incubated overnight at 4°C in the primary antibody. The IHC experiments noted include the use of chicken anti-GFP (Aves labs Cat#GFP-1020) and mouse anti-ChAT (Atlas Cat#AMAb91130). After primary incubation, sections were washed three times in 1xPBS and incubated in the same blocking solution containing secondary antibodies. Secondary antibodies used include goat anti-chicken IgY Alexa 488 (ThermoFisher Cat#A-11039), goat anti-mouse IgG2b Alexa 488 (ThermoFisher Cat#A-21141) and goat anti-mouse IgG2b Alexa 555 (ThermoFisher Cat#A-21147). Following secondary incubation, sections were washed three times and mounted.

For visualization of the retinas, whole eyes were removed, retinas were dissected, cut at the sides, and laid flat on a Superfrost slide fashioned with a Secure-Seal Spacer (ThermoFisher Cat#S24737), such that the photoreceptors were facing up.

To visualize direct SYFP2, direct EYFP, direct tdTomato, anti-Chat, and anti-GFP fluorescence, sections were mounted onto Superfrost slides (Fisher Cat#12-550-15) using VECTASHIELD Hardset antifade mounting medium with DAPI (Vector Laboratories Cat#H-1500-10) and imaged on a Nikon Eclipse Ti2 epifluorescence microscope, a Zeiss LSM 880 confocal microscope, or Leica SP8 confocal microscope using the manufacturer’s software.

#### Rat ICV injections of AAVs, immunohistochemistry, and imaging

Rat neonatal pups P0-P1 were injected using the following coordinates to hit the lateral ventricle: 1.5 mm lateral to bregma, 1.5 mm posterior to bregma, and 2 mm deep. A total of ∼1.50-2.00 × 10^11^ GC was delivered in 5µL total volume per hemisphere, and the needle was held in place in the ventricle for 30 seconds to allow virus diffusion prior to removal. Injected animals were returned to their home cage with the dam and euthanized at P18-P19 for analysis of viral expression.

For SYFP2 and cholinergic positive neuron detection, fixed sections were washed three times in 1xPBS at room temperature with gentle shaking, blocked in 1xPBS containing 5% normal goat serum (Vector Laboratories Cat#S-1000-20), 0.2% Triton X-100 (Millipore Sigma Cat#X100), bovine serum albumin (Millipore Sigma Cat#A9418-5G) for up to 2 hours at room temperature with gentle shaking, and then incubated overnight at 4°C in the primary antibody. The IHC experiments noted include the use of mouse anti-ChAT (Atlas Cat#AMAb91130). After primary incubation, sections were washed three times in 1xPBS and incubated in the same blocking solution containing secondary antibodies. Secondary antibodies used include goat anti-mouse IgG2b Alexa 555 (ThermoFisher Cat#A-21147).

Following secondary incubation, sections were washed three times and mounted with VECTASHIELD Hardset antifade mounting medium with DAPI (Vector Laboratories Cat#H-1500-10). Images were taken on a Nikon Eclipse Ti2 epifluorescence microscope or a Zeiss LSM 880 confocal microscope, using the manufacturer’s software.

#### Intra-cisterna magna (ICM) and intrathecal (IT) injections of AAV vectors into macaque, histology, immunochemistry, and imaging

For injections into macaque, large scale purified viral vector preps were packaged of the PHP.eB serotype by a commercial source (Packgene, Houston TX). The average titer of these viral preps was 9.0 × 10^13^ GC/ml. One week prior to survival surgery, an MRI scan was obtained for each animal and used to establish the injection coordinates.

For the IT injection, a three-year-old male macaque weighing 4.53 kg was anesthetized, placed on their side with their hind limbs towards the umbilicus such that there was a widening of the intervertebral space. A 22-gauge spinal needle was inserted into the L3-L4 intervertebral space, and its position was confirmed by CSF return. The syringe containing the virus was attached to the spinal needle and virus was injected using a syringe pump (Medfusion 2001) at a speed of 0.03mL per minute, with 2.27mL total volume injected at 1.0 × 10^13^ vg/kg. The tubing attached to the syringe was backfilled with saline and flushed immediately after to account for the dead space within the tubing.

For the ICM injections, two to three-year-old male and female macaques weighing 3-4 kg were anesthetized and placed in ventral recumbency with their head ventroflexed. A 22-gauge spinal needle was inserted into the area between the occipital protuberance and the C1 vertebrae and its position was confirmed by CSF return. The syringe containing the virus was attached to the spinal needle and virus was injected using a syringe pump (Medfusion 2001) at a speed of 0.03mL per minute, with 2.38mL total volume injected at 1.0 × 10^13^ vg/kg. The tubing attached to the syringe was backfilled with saline and flushed immediately after to account for the dead space within the tubing.

Animals were euthanized 5-7 weeks post-injection and the spinal cord, brain, and several other peripheral organs were collected. Fresh tissue was drop-fixed in 10% formalin for 48 hours at 4°C and then transferred to a 30% sucrose solution and kept at 4°C for several days to cryoprotect. For visualization of macaque heart, liver, kidney, pancreas, gastrocnemius muscle, and spinal cord (cervical, thoracic, lumbar, sacral), transverse sections were cut from each organ at 30µm and mounted onto Superfrost slides (Fisher Cat#12-550-15). Coronal brain sections were cut at 30µm and mounted onto Brain Research Laboratories slides (Cat#5075-W). All tissues were cut using a freezing microtome (Leica SM2000R).

For Flag-mTFP1 and cholinergic positive neuron detection, fixed sections were washed three times in 1xPBS at room temperature with gentle shaking, blocked in 1xPBS containing 5% normal goat serum (Vector Laboratories Cat#S-1000-20), 0.2% Triton X-100 (Millipore Sigma Cat#X100), bovine serum albumin (Millipore Sigma Cat#A9418-5G) for up to 2 hours at room temperature with gentle shaking, and then incubated overnight at 4°C in the primary antibody. The IHC experiments noted include the use of chicken anti-GFP (Aves labs Cat#GFP-1020), mouse anti-FLAG M2 (Sigma Cat#F1804) and mouse anti-ChAT (Atlas Cat#AMAb91130). After primary incubation, sections were washed three times in 1xPBS and incubated in the same blocking solution containing secondary antibodies. Secondary antibodies used include goat anti-chicken IgY Alexa 488 (ThermoFisher Cat#A-11039), goat anti-mouse IgG1 Alexa 488 (ThermoFisher Cat#A-21121), and goat anti-mouse IgG2b Alexa 555 (ThermoFisher Cat#A-21147).

Following secondary incubation, sections were washed three times and mounted with Vectashield Antifade Mounting Medium with DAPI (Vector Labs Cat#H-1200-10) or ProLong Gold Antifade mounting medium (Invitrogen Cat#P36930. Images were acquired on a Leica SP8 confocal microscope or a Nikon Eclipse Ti2 epifluorescence microscope using the manufacturer’s software.

#### RNAscope on mouse spinal cord sections and analysis

We performed RO injections of wild-type C57BL/6J mice with enhancer AAVs driving SYFP2, sacrificed animals 4-5 weeks post-injection and harvested the spinal cord tissue. Following dissection, spinal cords were embedded in optimum cutting temperature compound (OCT; Sakura TissueTek Cat#4583) and then stored at -80°C until ready to section. Immediately prior to sectioning, the spinal cords were thawed for one hour at –20°C and then cut at a thickness of 20µm on a cryostat. Thin sections were collected directly onto SuperFrost slides (ThermoFisher Scientific Cat#J3800AMNZ) and then left to dry for up to one hour at -20°C to ensure adhesion of the section to the slide. Slides were kept at –80°C for up to two weeks before single molecule fluorescent *in situ* hybridization was performed using the RNAscope Fluorescent Multiplex Reagent Kit V2 (Advanced Cell Diagnostics Cat#323100). The assay was carried out using the manufacturer’s instructions for fresh frozen tissue apart from one small deviation which was to fix the tissue for only 15 minutes in 4% PFA after removing slides from freezer. Probes used were *Chat* (ACD Cat#408731-C2) to label LMNs broadly, *Chodl* (ACD Cat#450211-C3), and *Ret* (ACD Cat#431791-C3) to differentiate between LMN subtypes. Since native SYFP2 fluorescence degrades in fresh frozen sections, a probe against *SYFP2* mRNA (ACD Cat#590291) was used to identify cells labeled by the specified enhancer viruses. All sections were stained with DAPI and mounted with ProLong Gold Antifade mounting medium (Invitrogen Cat#P36930) per the RNAscope kit protocol.

Sections were imaged using a 10x objective on a Leica SP8 confocal microscope with resonance scanning. Approximately 20µm stacks with 1µm steps were collected and processed in LASX (Leica) software to produce maximum intensity projections for each channel.

#### Clearing and light sheet imaging of mouse spinal cord

Paraformaldehyde-fixed samples were preserved using SHIELD reagents (LifeCanvas Technologies) using the manufacturer’s instructions ^46^. Mouse samples were delipidated using LifeCanvas Technologies Clear+ delipidation reagents and rat samples were delipidated using LifeCanvas Technologies SmartBatch+ device for 2 days. Following delipidation, samples were incubated in 50% EasyIndex (RI = 1.52, LifeCanvas Technologies) overnight at 37°C followed by 1 day incubation in 100% EasyIndex for refractive index matching. After index matching, the samples were imaged using a SmartSPIM axially swept light sheet microscope using a 3.6x objective (0.2 NA) (LifeCanvas Technologies).

#### Western Blot

Cervical spine tissue from mice was flash-frozen on dry ice and stored at -80°C until processing. Samples were sonicated in 95°C 1% SDS with protease inhibitors (ThermoFisher Cat#78441), using previously described methods ^45^. Protein concentration was measured using a BCA assay kit (ThermoFisher Cat#23225), then denatured in NuPage LDS Sample Buffer (Cat#NP0007) and boiled at 95°C for 5 min. 75 µg of protein was separated on a 10–20% Tris/Glycine gel (ThermoFisher Cat#XP10202BOX) and transferred to a nitrocellulose membrane (ThermoFisher Cat#88025). Blots were incubated overnight with anti-GFP (Abcam Cat#ab6556, 1:2500) and β-actin (Millipore Cat#MAB1501R, 1:1000), followed by fluorescent secondary antibodies.

Detection was performed using an Odyssey Infrared Imaging System (LI-COR Biosciences). SYFP (GFP antibody signal) was normalized to β-actin and then to the average HCT1 levels for each blot.

### Quantification and statistical analysis

Cell counts of epifluorescence or confocal images were obtained using the ImageJ analysis Cell Counter plug in and automatic spot-detection algorithm (Imaris 10.0.1, Bitplane AG). Briefly, the multichannel tiff files were uploaded into ImageJ, split into the individual channels, and brightness and contrast were adjusted uniformly across the images such that the signal could be easily distinguished from the background. The individual channel images were then compressed into a stack and the number of cells visible in each channel were counted. For Imaris, the images were converted into ims files using plug in, brightness and contrast kept uniform across all, spots were identified for individual channels using the automated algorithm (15µm), and colocalization was assigned automatically with a constant shortest distance to spots (<10µm).

For IHC and double transgenic experiments, percent specificity for a given virus was calculated by taking the number of double positive cells (i.e., SYFP2+ and Chat+ or tdTomato+) and dividing that number by the total number of virally labeled cells (i.e. SYFP2+). Percent completeness of labeling was calculated by taking the number of double positive cells (i.e., SYFP2+ and Chat+ or tdTomato+) and dividing that by the total number of positive cells for the cell type of interest population (i.e., Chat+ or tdTomato+).

For RNAscope experiments, we quantified specificity and completeness of our viruses against subtype selective probes, *Ret* and *Chodl*. Percent specificity for a given virus was calculated by taking the number of double positive cells (i.e., *SYFP2*+ and *Ret*+ or *Chodl*+) and dividing that number by the total number of virally labeled cells (i.e., *SYFP2*+). Percent completeness of labeling was calculated by taking the number of double positive cells (i.e., *SYFP2*+ and *Ret*+ or *Chodl*+) and dividing that by the total number of probe-labeled cells (i.e., *Ret*+ or *Chodl*+).

To validate the probes used in RNAscope experiments, we quantified specificity and completeness of the probes against the pan-LMN marker, *Chat,* as well as the average soma size of double positive cells (*Chat+* and *Ret+* or *Chodl+*). Percent specificity for the subtype selective probes was calculated by taking the double positive cells (i.e., *Ret*+ or *Chodl*+ and *Chat*+) and dividing that number by the total number of *Ret*+ or *Chodl*+ cells. Percent completeness of labeling was calculated by taking the number of double positive cells (i.e., *Ret*+ or *Chodl*+ and *Chat*+) and dividing that by the total number of *Chat*+ positive cells. Cell sizes were collected using Measurement Points in Imaris with the sphere/pair option where the distance was automatically calculated with two points (the longer diameter) of a *Chat+* cell that colocalized with *Ret* or *Chodl*.

Values were imported, analyzed, and graphed using GraphPad Prism 10 software (Dotmatics, Boston, MA) and data are presented heatmaps or bar graphs as mean ± standard error from the mean. Statistical comparisons of means within the same treatment group was performed using a one-way ANOVA or using a two-way ANOVA for comparison of the same virus injected via different routes of administration.

## Supplemental Information

**Supplemental Figure 1.**
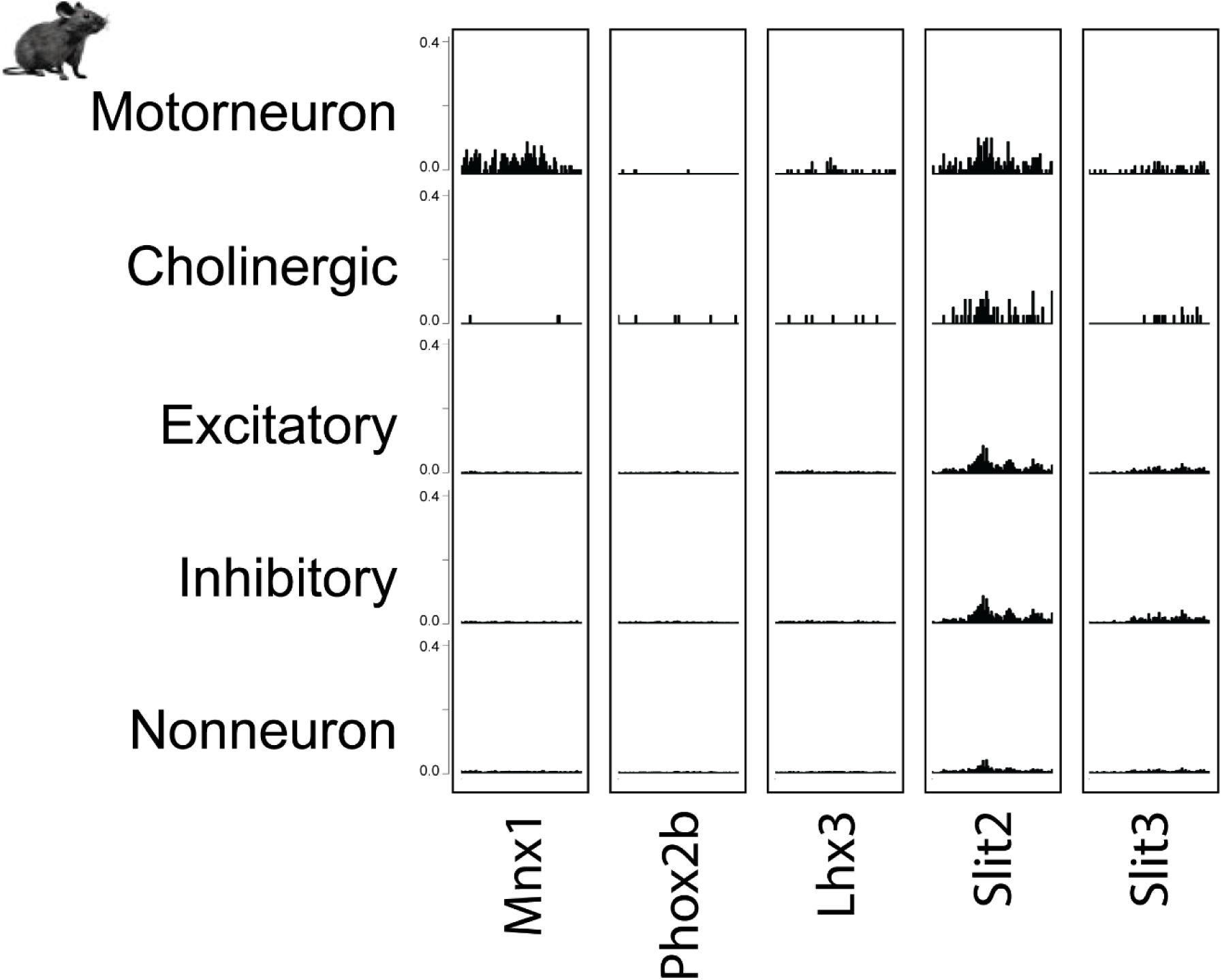
Chromatin accessibility of regulatory regions for reported motor neuron marker genes. Chromatin accessibility tracks for each major cell type from mouse ATAC-seq. The accessibility is measured within the regulatory region (+/- 2.5kb of TSS) for previously reported motor neuron marker genes.

**Supplemental Figure 2.**
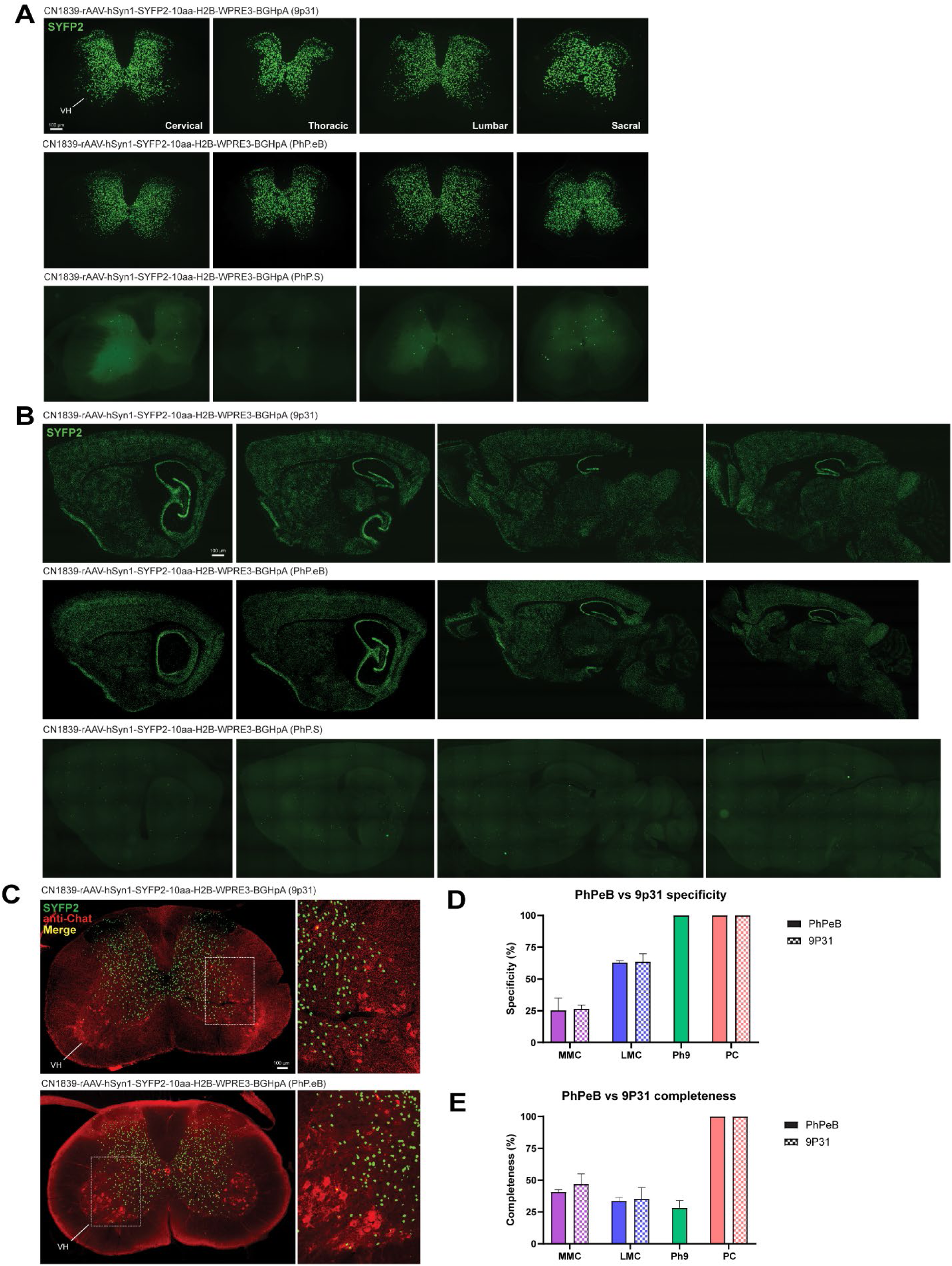
Capsid screening in mouse spinal cord and brain. (**A-B**) AAVs containing the pan neuronal hSyn1 promoter driving SYFP2 fluorescent protein expression were packaged with three candidate BBB permeant capsids (PHP.eB, 9P31, PHP.S) and tested by RO injection at a dose of 5.0 × 10^11^ GC in wild-type mice. Representative epifluorescence images of native SYFP2 fluorescence in 50µm thick transverse sections from each level of cord (**A**) or 25µm thick sagittal brain sections (**B**) are shown for all CN1839 viruses. (**C**) Representative epifluorescence images of CN1839-driven native and nuclear-enriched SYFP2 fluorescence (green) and anti-Chat staining (red) in 30µm thick cervical spinal cord sections. Higher magnification images of cells boxed in the low magnification images are presented in the right panels. (**D-E**) Quantification of percent (%) specificity and completeness of PHP.eB CN1839 vs 9P31 CN1839 in anatomical areas of interest or in partition cells within cervical spinal cord. Please refer to main Figure 2 to see region and cell type labels at a similar anatomical level. Data are presented as mean ± S.E.M for n=2 animals and 3 sections per animal analyzed.

**Supplemental Figure 3.**
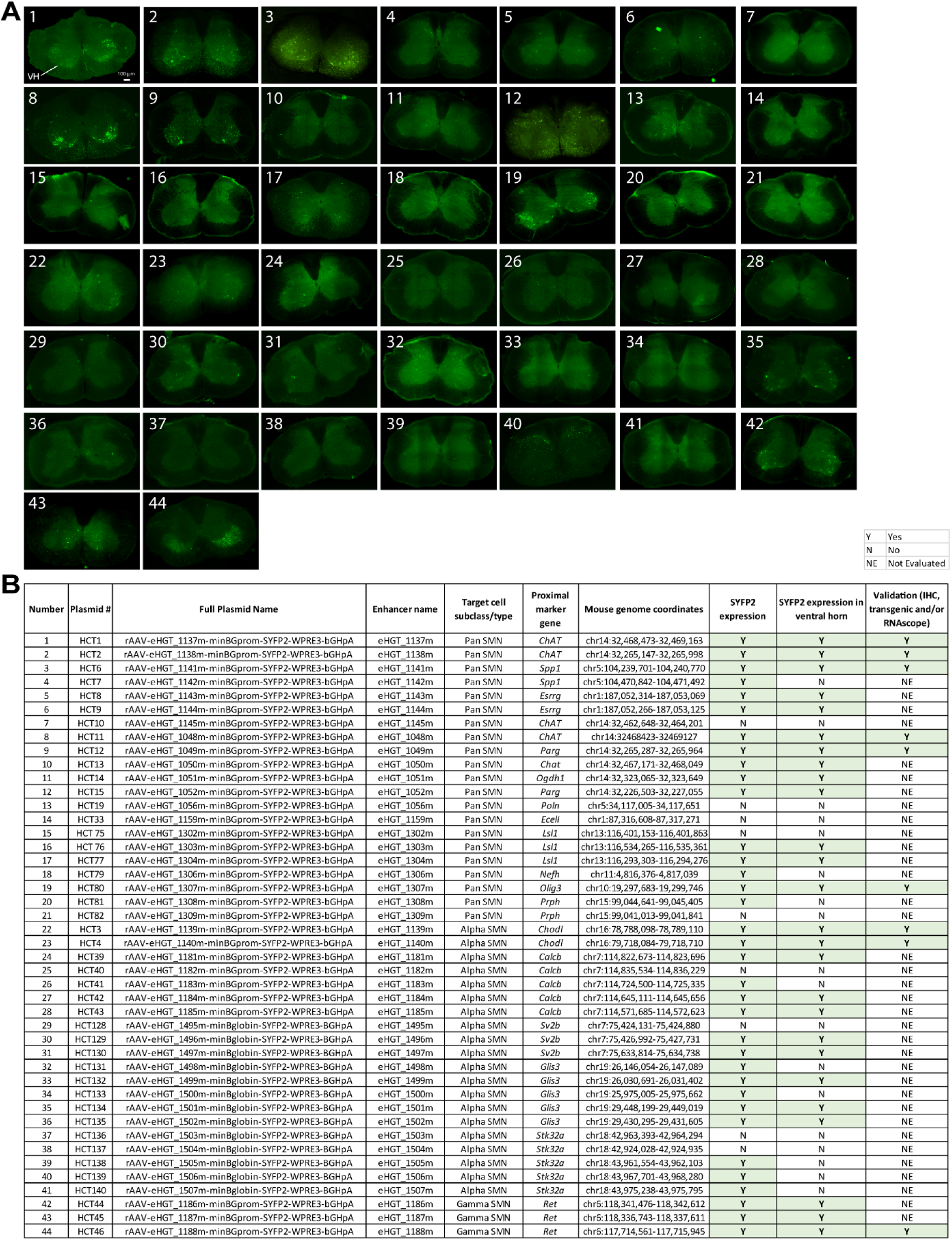
Initial screening of putative pan, alpha, and gamma spinal motor neuron enhancer viruses in mouse, related to Figure 2. (**A**) Candidate enhancers were functionally screened in C57Bl/6J mice by RO injection of SYFP2-expressing viruses. Representative epifluorescence images of native SYFP2 fluorescence in 50µm thick transverse cervical sections are shown for all viruses screened. VH = ventral horn and number corresponds to the summary table in panel **B**. Enhancer viruses with no visible cells labelled throughout the sections analyzed = No (N) and those with some positive SYFP2 cells in any location of the section = Yes (Y) in SYFP2 expression category column. Viruses labeling cells specifically within the ventral horn = Y in SYFP2 expression in ventral horn column. Promising viruses that were selected for additional molecular validation are denoted as Y in the the last column.

**Supplemental Figure 4.**
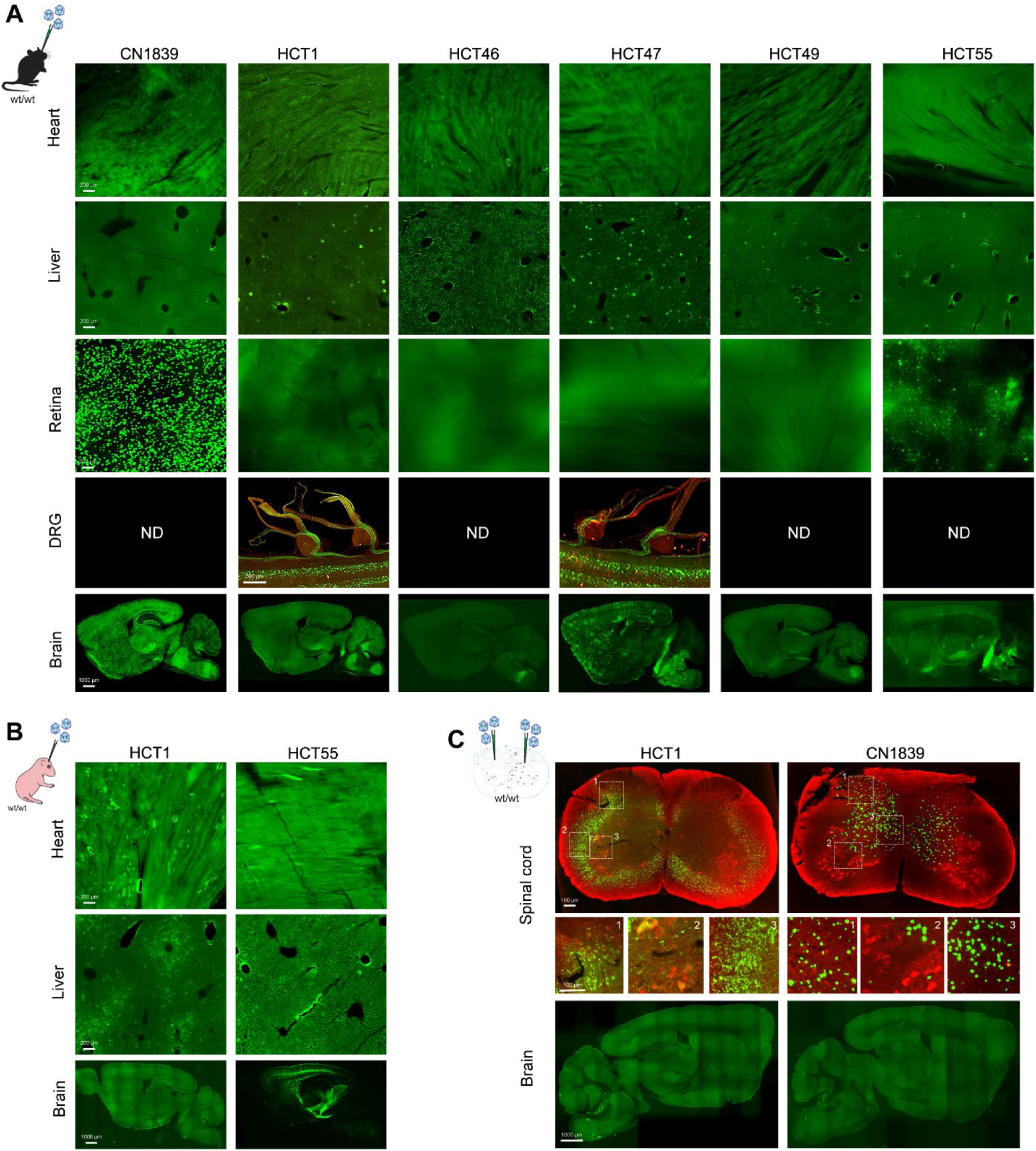
CNS and peripheral organ biodistribution data from mice administered AAVs via different routes. (**A**) Representative epifluorescence images of native SYFP2 fluorescence in heart (50µm), liver (30µm), retina (whole mount), DRG (dorsal root ganglion; from 3D light sheet image of intact cord) and brain (25µm) tissue following RO injection of the indicated viruses. (**B**) Representative epifluorescence images of native SYFP2 fluorescence in heart (50µm), liver (30µm) and brain (25µm) tissue following ICV injection of mouse neonatal pups. (**C**) Representative epifluorescence images of HCT1- or CN1839-driven native SYFP2 fluorescence (green) and anti-Chat signal (red) in mouse transverse spinal cord sections (30µm) or brain sections (25µm) following bilateral intraspinal injection (IS) of 1.0 × 10^13^ GC; 250nL left hemisphere and 50nL right hemisphere.

**Supplemental Figure 5.**
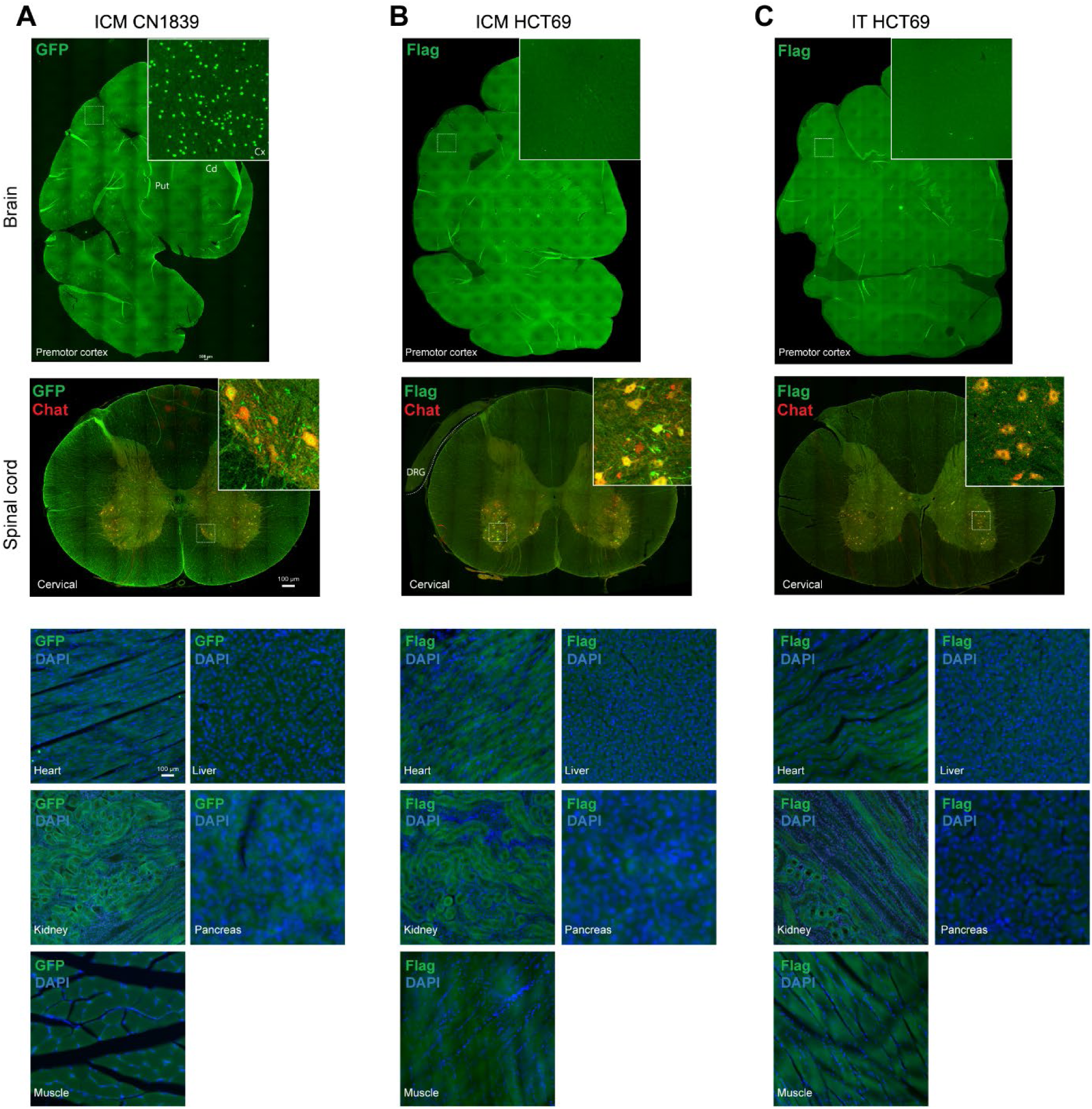
CNS and peripheral organ biodistribution data for HCT69 and CN1839 viruses following ICM or IT routes of administration to macaque. Representative epifluorescence images of anti-Flag signal (green) or anti-GFP signal (green) in coronal brain sections prepared from CN1839 ICM (**A**) HCT69 ICM (**B**) or HCT69 IT (**C**) PHP.eB virus injected animals. Representative epifluorescence images of anti-Flag signal (green) or anti-GFP signal (green) merged with anti-Chat signal (red) in transverse spinal cord sections prepared from CN1839 ICM (**A**) HCT69 ICM (**B**) or HCT69 IT (**C**) PHP.eB virus injected animals (DRG, dorsal root ganglion). Representative epifluorescence images of anti-Flag signal (green) or anti-GFP signal (green) merged with anti-DAPI signal (blue) in transverse peripheral tissue sections prepared from CN1839 ICM (**A**) HCT69 ICM (**B**) or HCT69 IT (**C**) PHP.eB virus injected animals.

**Supplemental Figure 6.**
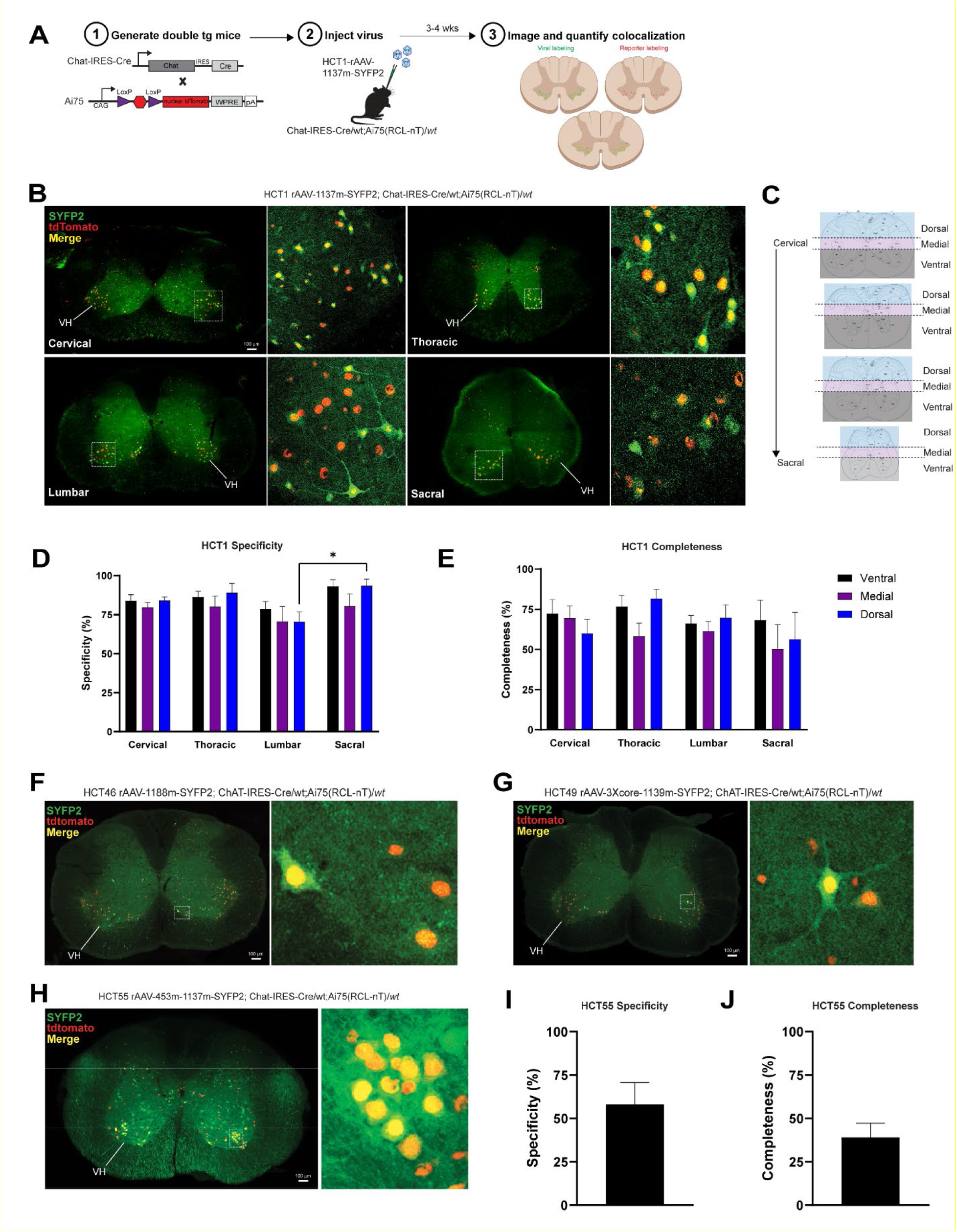
Additional validation of enhancer AAVs in double transgenic mice. (**A**) Experimental workflow for testing the indicated viruses using ChAT-IRES-Cre;Ai75 mice. (**B**) Representative confocal images of RO-injected, HCT1-driven native SYFP2 (green) fluorescence and nuclear tdTomato (red) fluorescence in 50µm thick transverse spinal cord sections from double transgenic mice. (**C**) Schematic of coarse segmentation plan for quantification of viral labeling in transverse spinal cord sections. Colored areas denote segmentation overlayed onto reference images from the Allen Institute mouse spinal cord atlas. (**D-E**) Quantification of HCT1 virus specificity and completeness of labeling throughout all segments and levels of cord. Data are plotted as mean ± SEM and *p = 0.0253 (n = 3 animals and 3 sections per level per animal analyzed). (**F-H**) Representative confocal images of native SYFP2 (green) fluorescence and nuclear tdTomato (red) fluorescence in 50µm transverse spinal cord sections from (**F**) HCT46-, (**G**) HCT49-, and (**H**) HCT55-injected double transgenic mice. (**I**) Quantification of HCT55 virus specificity and completeness of labeling in cervical cord. Data are plotted as mean ± SEM and *p < 0.05 (n = 2 animals and 3 sections for each level per animal analyzed). VH = ventral horn and cells bounded by white boxes in the low magnification images are presented in the high magnification images to the right of each.

**Supplemental Figure 7.**
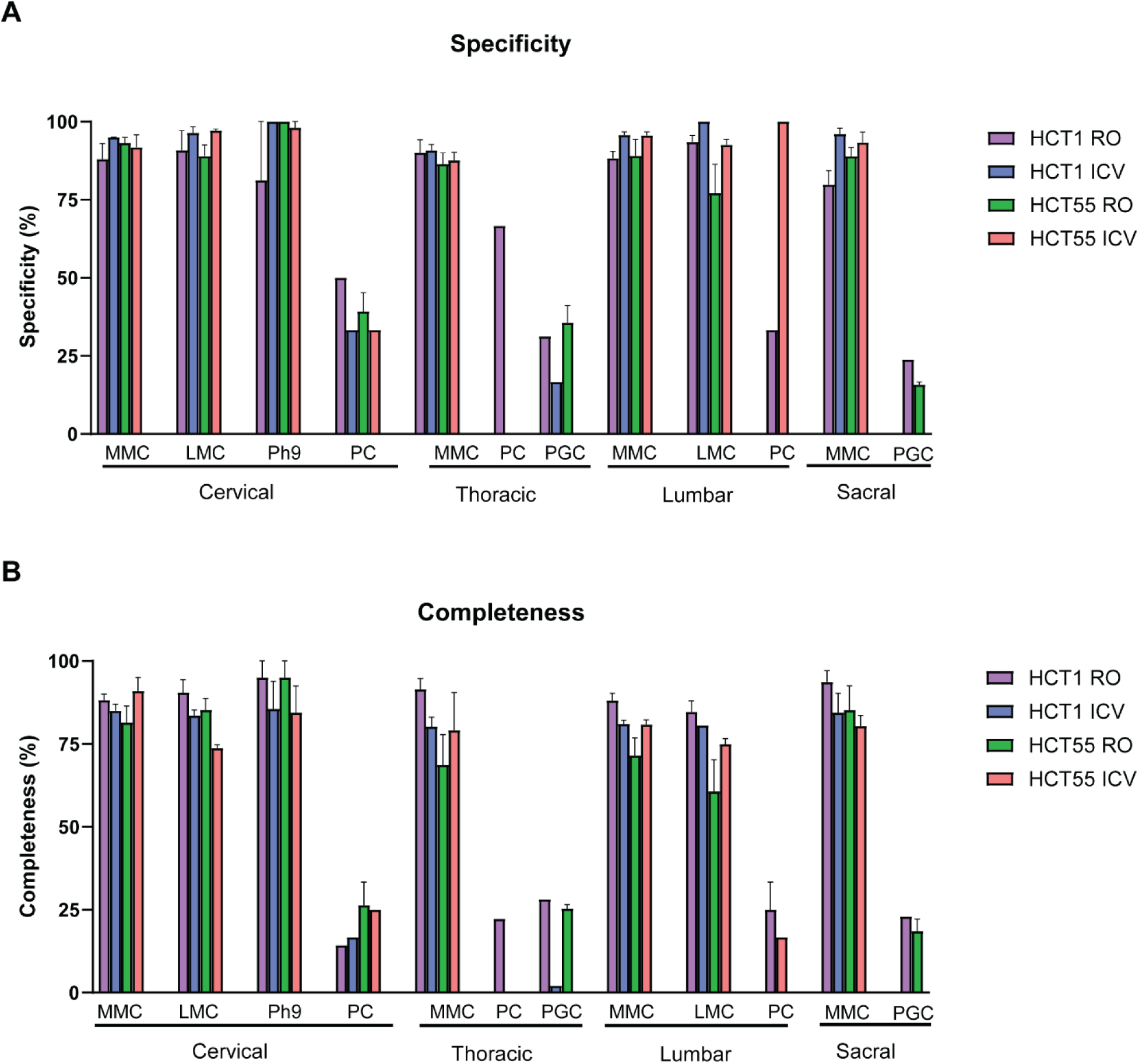
Summary plots to track variance. Quantification of specificity (**A**) and completeness (**B**) of HCT1 or HCT55 viral labeling of Chat+ motor neurons as percent (%) in the different levels of spinal cord following RO or ICV injection. Anatomical areas of interest or previously described cell types are denoted and labelled below the bars. MMC = medial motor column, LMC = lateral motor column, Ph9 = phrenic motor neurons of lamina 9, PGC = preganglionic motor column, and PC = partition cells. Data are plotted as mean ± SEM and *p < 0.05 (n = 3 animals and 3 sections for each level per animal analyzed). The absence of an error bar denotes the conditions where either the anatomical areas/the cell type could be identified or colocalization of viral labeling with Chat+ motor neurons were present in only 1 animal out of the 3 analyzed.

**Table S1. Predicted accessibility of all putative pan LMN enhancers.** Accessibility of putative pan LMN enhancers in human whole body ATAC-Seq ^21^ and whole mouse brain ATAC-Seq ^22^. For each enhancer we report the summed accessibility within the enhancer’s genomic region for each cell type or tissue.

**Table S2. Predicted transcription factor motifs within the 1137m enhancer sequence.** Enrichment of transcription factor motifs from the JASPAR CORE database within the DNA sequence of 1137m.

**Table S3. Summary table of AAV biodistribution data.** AAV, adeno-associated virus; ICM, intra-cisterna magna; IT, intrathecal; IS, intraspinal; RO, retro-orbital; ICV, intracerebroventricular; DRG, dorsal root ganglion. - = no expression, + is 0 to 5 cells, ++ is 6 to 30 cells, +++ is 30 to100 cells, and ND is not done. HCT1 is pan LMN eHGT 1137m SYFP2 virus, HCT46 is gamma LMN eHGT 1188m SYFP2 virus, HCT47 is pan LMN 3Xcore eHGT 1137m virus, HCT49 is alpha LMN eHGT 1139m SYFP2 virus, HCT55 is a dual stitched L5 ET and pan LMN, eHGT453m and eHGT 1137m SYFP2 virus, HCT69 is pan LMN 1137m-3XFLAG-mTFP1 virus and CN1839 is pan neuronal H2B-SYFP2 virus.

**Video S1. Light sheet video of an HCT1 virus infected mouse spinal cord**. Native SYFP2 fluorescence (green) is shown throughout intact spinal cord and in transverse spinal cord sections.

**Video S2. Light sheet video of an CN4358 virus infected mouse spinal cord**. Native SYFP2 fluorescence (green) is shown throughout intact spinal cord and in transverse spinal cord sections.

**Video S3. Light sheet video of an HCT1 virus infected rat spinal cord**. Native SYFP2 fluorescence (green) is shown throughout intact spinal cord and in transverse spinal cord sections.

**Video S4. Light sheet video of an CN1839 virus infected rat spinal cord**. Native SYFP2 fluorescence (green) is shown throughout intact spinal cord and in transverse spinal cord sections.

**Video S5. Light sheet video of an HCT47 virus infected mouse spinal cord**. Native SYFP2 fluorescence (green) is shown throughout intact spinal cord and in transverse spinal cord sections.

