## Supplemental Table 3 for "Enhancer AAVs for targeting spinal motor neurons and descending motor pathways in rodents and macaque"

**Table S3.** Summary table of AAV biodistribution data (Related to Figure S4 and S5).

| Species | Mouse | Mouse | Mouse | Mouse | Mouse | Mouse | Mouse | Mouse | Macaque | Macaque | Macaque | Mouse | Mouse |
|---|---|---|---|---|---|---|---|---|---|---|---|---|---|
| Injection | IS | RO | ICV | RO | RO | RO | RO | ICV | ICM | IT | ICM | IS | RO |
| Virus | HCT1 | | | HCT46 | HCT47 | HCT49 | HCT55 | | HCT69 | | CN1839 | | |
| Cell type target | Pan-LMN | | | Gamma-LMN | Pan-LMN | Alpha-LMN | L5 ET + pan-LMN | | Pan-LMN | | Pan-neuronal | | |
| Brain | + | + | +++ | + | +++ | + | +++ | +++ | - | - | +++ | + | +++++ |
| Spinal cord | + | +++ | +++ | +++ | ++++ | +++ | ++++ | ++++ | ++ | ++ | + | +++++ | +++++ |
| DRG | ND | - | ND | ND | - | ND | ND | ND | - | ND | ND | ND | ND |
| Heart | ND | - | - | - | - | - | - | - | - | - | + | ND | - |
| Liver | ND | + | - | - | + | - | - | - | - | - | - | ND | - |
| Kidney | ND | ND | ND | ND | ND | ND | ND | ND | - | - | - | ND | ND |
| Pancreas | ND | ND | ND | ND | ND | ND | ND | ND | - | - | - | ND | ND |
| Skeletal muscle | ND | ND | ND | ND | ND | ND | ND | ND | - | - | - | ND | ND |
| Retina | ND | - | ND | - | - | - | + | ND | ND | ND | ND | ND | +++++ |
